# An unrecognized host response to microbial exposure resets circadian timing

**DOI:** 10.64898/2026.04.11.717924

**Authors:** Devons Mo, Ti (Dung Ngoc) Lam, Ella Baker, Olivia P. Fraser, Jack Dorling, John S. O’Neill, Gerben van Ooijen, Antony N. Dodd, Carrie L. Partch, Priya Crosby, Jacqueline M. Kimmey

**Affiliations:** Department of Microbiology and Environmental Toxicology, University of California, Santa Cruz; Santa Cruz, USA; Department of Cell and Developmental Biology, John Innes Centre; Norwich Research Park, Norwich, UK; Institute of Molecular Plant Sciences, School of Biological Sciences, University of Edinburgh; Edinburgh, UK; MRC Laboratory of Molecular Biology; Cambridge, UK; Department of Chemistry and Biochemistry, University of California, Santa Cruz; Santa Cruz, USA; Howard Hughes Medical Institute, University of California, Santa Cruz; Santa Cruz, CA USA; Center for Circadian Biology, University of California, San Diego; La Jolla, CA USA

## Abstract

As ubiquitous features of every natural environment, microbes have profoundly shaped eukaryotic biology throughout evolution. Circadian clocks evolved in all domains of life as central regulators that align physiology with environmental cycles, yet whether they respond directly to microbial signals remains unknown. Here, we demonstrate that evolutionarily diverse microbes potently reset mammalian cellular clocks and can drive phase shifts in plants and algae, indicating cross-kingdom effects of microbes on circadian rhythms. In mammals, exposure to soluble bacterial components distinct from canonical innate immune ligands induces acute PER2 upregulation independently of *Bmal1* or nascent transcription. A targeted inhibitor screen and biochemical assays implicate p38 MAPK as a modulator of this response. Taken together, this positions bacterial exposure as a previously unrecognized circadian clock input, revealing a new axis of host-microbe interaction that modulates biological timing at the cellular level.

## Introduction

Microorganisms are ubiquitous and evolutionarily enduring life forms that inhabit every environment on Earth. Approximately 2 billion years ago, microbial symbiosis enabled the formation of eukaryotic cells, allowing complex life to evolve^1^. Today, microbes continue to influence core biological processes including immunity^2^, metabolism^3^, and behavior^4^, yet microbial influence is often overlooked in biological models. Only in the past two decades, for example, have we begun to appreciate how the human microbiome shapes health and physiology. We now understand that products such as bacterial cell wall and metabolites influence diverse processes including epithelial turnover, nutrient absorption, and systemic immune tone^2,3,5^. These signals fluctuate with microbial composition and activity, which oscillate based on environmental changes in the host such as feeding patterns and hormonal states. Through these dynamics, microbial inputs contribute to rhythmic processes in host tissues, including gene expression, barrier function, and immune surveillance^2,5,6^.

Microbial communities across ecosystems undergo robust daily oscillations in both composition and biomass shaped by changes in light, temperature, and other environmental signals^7–12^. Even the air we breathe is rhythmic, with microbial biomass rising several-fold at night and bacterial versus fungal taxa alternating in dominance across the day^8,13^. In soil, the rhizosphere microbiome also undergoes diurnal restructuring, partly influenced by plant circadian rhythms^10^. In parallel, roughly 15% of gut taxa oscillate daily^14^, driven by host behaviors such as feeding and circadian rhythms. Together, these examples underscore that microbes are temporally dynamic, raising the provocative possibility that microbial signals themselves act as zeitgebers for eukaryotic circadian clocks.

Circadian clocks are cell-autonomous oscillators that evolved to align physiology with environmental cycles. In mammals, the core circadian clock is driven by the heterodimeric transcription factor CLOCK:BMAL1, which drives expression of its repressors, PER and CRY, to generate a ∼24-hr transcriptional translational feedback loop (TTFL)^15^. To stay aligned with Earth’s 24-hour cycle, circadian clocks synchronize to recurring environmental cues such as light^16^, temperature^17^, and feeding^18^. Despite widespread microbial influence on host biology^19–21^, it remains unknown whether microbial signals can act directly on the core circadian clock. Here, we address this question, revealing a previously unrecognized layer of communication between microbial signals and circadian timing.

## Results

### Circadian phase shifts are a common outcome of exposure to bacteria

To determine whether bacterial exposure directly affects the mammalian clock, we monitored the core circadian protein PER2 using mouse lung fibroblasts (MLF) derived from PERIOD2::LUCIFERASE (P2L) mice. This strain expresses luciferase fused to PER2 at the *Per2* endogenous locus, such that bioluminescence tracks PER2 protein abundance^22^. To enable multi-day recordings without bacterial overgrowth, all bacteria were heat-killed (HK). P2L MLFs were exposed to a panel of evolutionarily diverse bacteria and PER2 rhythms were monitored (experimental schematic, Fig. 1A). Addition of an equivalent volume of phosphate-buffered saline (PBS) served as our bacteria-free control. Surprisingly, in our initial screen, evolutionarily diverse, pathogenic Gram-positive and Gram-negative bacteria all shifted the circadian phase relative to PBS controls (Fig. 1B). Exposure to bacteria triggered an acute increase in PER2 evident within 1-hr that peaked between 4 to 6-hrs (Extended data Fig. 1A). We subsequently tested exposure to the pathogen *Listeria monocytogenes*, environmental organisms *Mycobacterium smegmatis* and *Bacillus subtilis*, and a non-pathogenic strain of *Escherichia coli* – all of which induced phase shifts in P2L MLF (Extended data Fig. 1B-C). Because responses might depend on circadian phase, we repeated exposures to *Streptococcus pneumoniae* (hereafter abbreviated as *Spn*), *Staphylococcus aureus* and *Escherichia coli.* These again shifted rhythms, leading to almost a complete inversion when applied near the PER2 trough (Fig. 1C).

**Figure 1.**
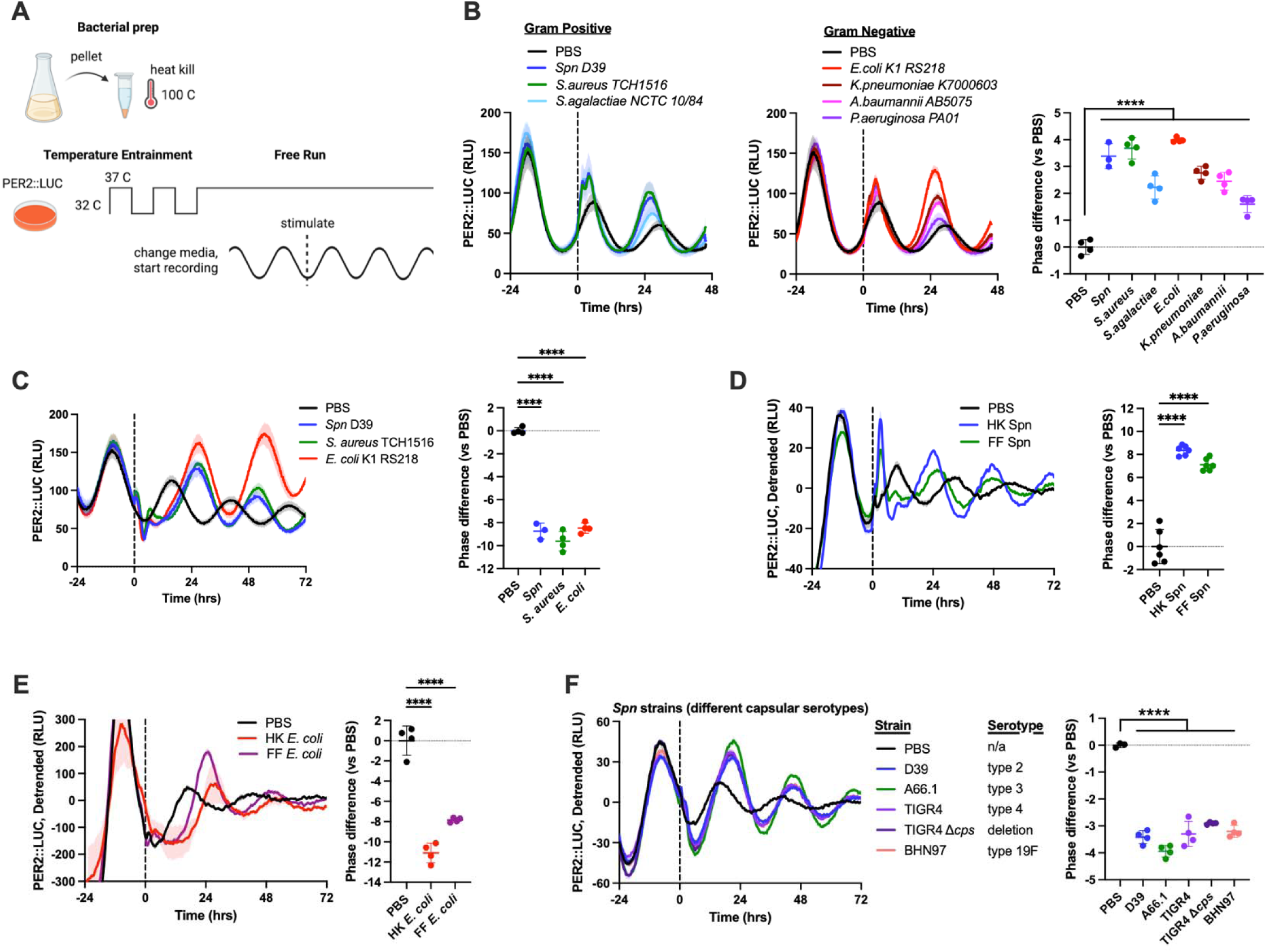
Circadian phase shifts are a common outcome of exposure to bacteria. (**A**) A diagram depicting the preparation of HK bacteria and the experimental procedure. (**B**) Relative luminescence units (RLU) were monitored from PER2::LUC mouse lung fibroblasts (P2L MLF) before and after stimulation with HK bacteria at time 0 (dashed line). The differences in phase between bacteria treated and PBS treated samples are calculated in circadian hours and graphed as “phase difference”. A negative value indicates a phase delay and a positive value indicates a phase advance. Circadian time (CT) is defined such that CT0/24 is the trough and CT12 is the peak of PER2 expression. Relative luminescence units (RLU). (**C**) Several bacterial species from (A) were retested to determine their ability to drive changes in PER2 phase. (**D-E**) P2L MLF were stimulated with formalin-fixed (FF) and HK *Spn* (D) or *E. coli* (E) and compared to PBS controls. **(F)** P2L MLF were stimulated with various strains of *Spn* which each express a different capsular serotype. Recording traces are shown as mean±S.E.; all other graphs are shown as mean±S.D. **** = p<0.0001. *n* ≥ *3* for all groups, dots indicate technical replicates. See Table S2 for all sample sizes and detailed statistical results.

To rule out the possibility that heat-inactivation introduced an artifact, we tested whether fixed bacteria also drive changes in PER2 phase. As shown in Fig. 1D-E, formalin-fixed (FF) *Spn* and *E. coli* induced similar phase shift responses as HK bacteria. Unlike heat-inactivation, which lyses cells and denatures proteins, formalin-fixation preserves cellular integrity^23^. Hence, the presence of PER2 phase shifts caused by FF bacteria suggests that at least some of the ligands responsible for inducing the phase shift response were accessible on the cell surface.

Many bacteria produce an extracellular polysaccharide capsule to avoid detection by the host^24^. Over 100 capsular serotypes of *Spn* with different sugar composition, linkage, and length have been identified, which alter host-bacterium interactions^25,26^. In addition to the previously tested *Spn* D39 strain (type 2 capsule), we tested A66.1 (type 3), TIGR4 (type 4), and BHN97 (type 19F) to determine if capsular serotype affects the phase shift response. D39 and TIGR4 have smaller capsules and cause invasive disease in mice, whereas A66.1 and BHN97 have larger capsules that lead to localized infections. We also used a capsule knockout (TIGR4 Δ*cps*) which is severely attenuated in virulence^27^. However, despite these marked differences in biological phenotypes, all tested strains of *Spn* induced equal phase shift responses (Fig. 1F).

### Initial detection of bacteria is sufficient to drive circadian resetting

To test whether closely spaced bacterial exposures produce additive effects on phase, we stimulated P2L MLFs with HK *E. coli* at one or both of two timepoints spaced 7 hours apart. Consistent with phase-dependent responsiveness, stimulation at time 0 produced a phase delay, whereas a single stimulation 7 hours later produced a phase advance. Dual stimulation resulted in a phase delay, similar to time 0 alone, indicating that the second stimulus did not produce an additional phase shift under these conditions (Fig. 2A). This result is consistent with the cellular sensing of various stimuli, as many physiological stimuli induce a period of temporary desensitization. For instance, the autophosphorylating insulin receptor loses kinase activity upon activation^28,29^, and many G-protein coupled receptors internalize into intracellular vesicles upon binding of their respective agonist, preventing further activation of downstream pathways ^30,31^. Importantly, both classes of receptors^32–34^ have been reported to transduce signals to the circadian clock, suggesting that desensitization to acute, repetitive stimuli does not necessarily exclude a given sensory pathway from a circadian role.

**Figure 2.**
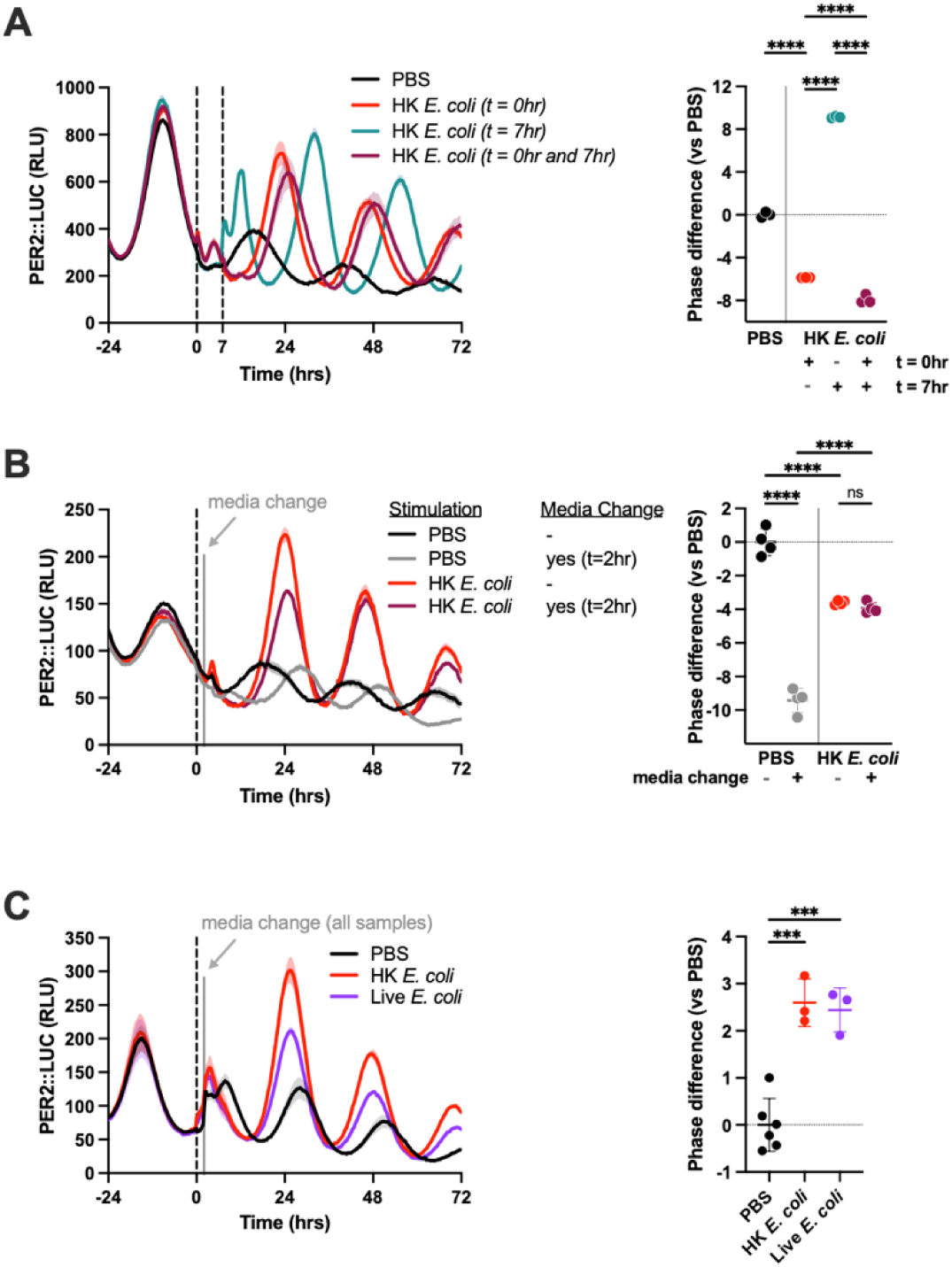
Transient bacterial exposure is sufficient to drive changes in PER2 phase. (**A**) P2L MLF were stimulated with HK *E. coli* at time 0 or 7 (dashed lines) or both. (**B**) P2L MLF were stimulated with HK *E. coli* or PBS (time 0, dashed line) and left untreated or were additionally washed at 2 hours post-stimulation (solid grey line). (**C**) P2L MLF were stimulated with live *E. coli* (time 0, dashed line) and washed at 2 hours post-stimulation (solid grey line) as compared to PBS controls. Recording traces are shown as mean±S.E.; all other graphs are shown as mean±S.D. *** = p<0.001; **** = p<0.0001; ns = not significant. *n* ≥ *3* for all groups, dots indicate technical replicates. See Table S2 for all sample sizes and detailed statistical results.

We next sought to test if continued bacterial exposure is necessary to drive changes in phase. We stimulated P2L MLFs with HK *E. coli* for two hours before removing the bacteria by washing with PBS then replacing the media. Exposure of cells to HK *E. coli* resulted in a ∼4 hour phase shift regardless of whether bacteria remained for the duration of the experiment or were removed two hours after stimulation, demonstrating that bacteria do not need to remain present to drive a phase shift (Fig. 2B). Consistent with known effects, media replacement alone shifted phase in PBS-treated controls. Interestingly, media exchange had no effect on cells exposed to bacteria, which mirrors the results obtained with dual bacterial stimulation (Fig. 2B-C).

Having established that sustained exposure to bacteria is not necessary for a circadian response and that washing does not abolish or obscure the HK *E. coli* effect, we next used this approach to test live bacteria, which must be removed to prevent rapid bacterial overgrowth that would interfere with circadian recordings. When live *E. coli* was added then removed two hours later, MLFs showed a significant phase shift closely mirroring the HK bacteria effect, showing that live bacteria were able to induce a circadian response as well (Fig. 2C).

### Bacterial signals reset circadian clocks across kingdoms

We next asked whether microbial stimulation shares other defining features of circadian input signals such as dose- and phase-dependent effects on the circadian clock^35^. As Gram-positive and Gram-negative bacterial lineages diverged over 2 billion years ago, we selected a member of each for further characterization^36^. *Spn* is a clinically relevant Gram-positive bacterium that colonizes the upper respiratory tract of humans and remains the leading cause of community-associated bacterial pneumonia^37^; the D39 strain remains highly virulent in murine models. In contrast, *E. coli* is Gram-negative, and DH5α is a lab-adapted, non-pathogenic strain that diverged from wild-type isolates through long-term culture since its isolation in 1922^38^.

First, we generated dose-response curves by stimulating P2L MLFs with varying doses of HK *Spn* or *E. coli* and measuring the resulting phase shifts. To prevent media-related artifacts, treatments were limited to 5% of the culture volume. Similar to previously-described resetting cues^35^, the magnitude of the phase shift that occurred in response to either bacterial species was dose-dependent (Fig. 3A-B). The largest shift occurred with 4-5×10^8^ colony forming units (CFU) / mL, quantified before heat-inactivation. We next stimulated cells at various circadian phases to generate phase response curves (PRC) and phase transition curves (PTC). The PRC confirmed that the shift magnitude depends on stimulation timing (Fig. 3C; double-plotted). Remarkably, the PTCs for both *Spn* and *E. coli* exhibited near-zero average slopes (Extended data Fig. 2A-B), indicating that bacterial exposure sets cells to approximately equal circadian times regardless of the initial phase – a feature of strong resetting cues.

**Figure 3.**
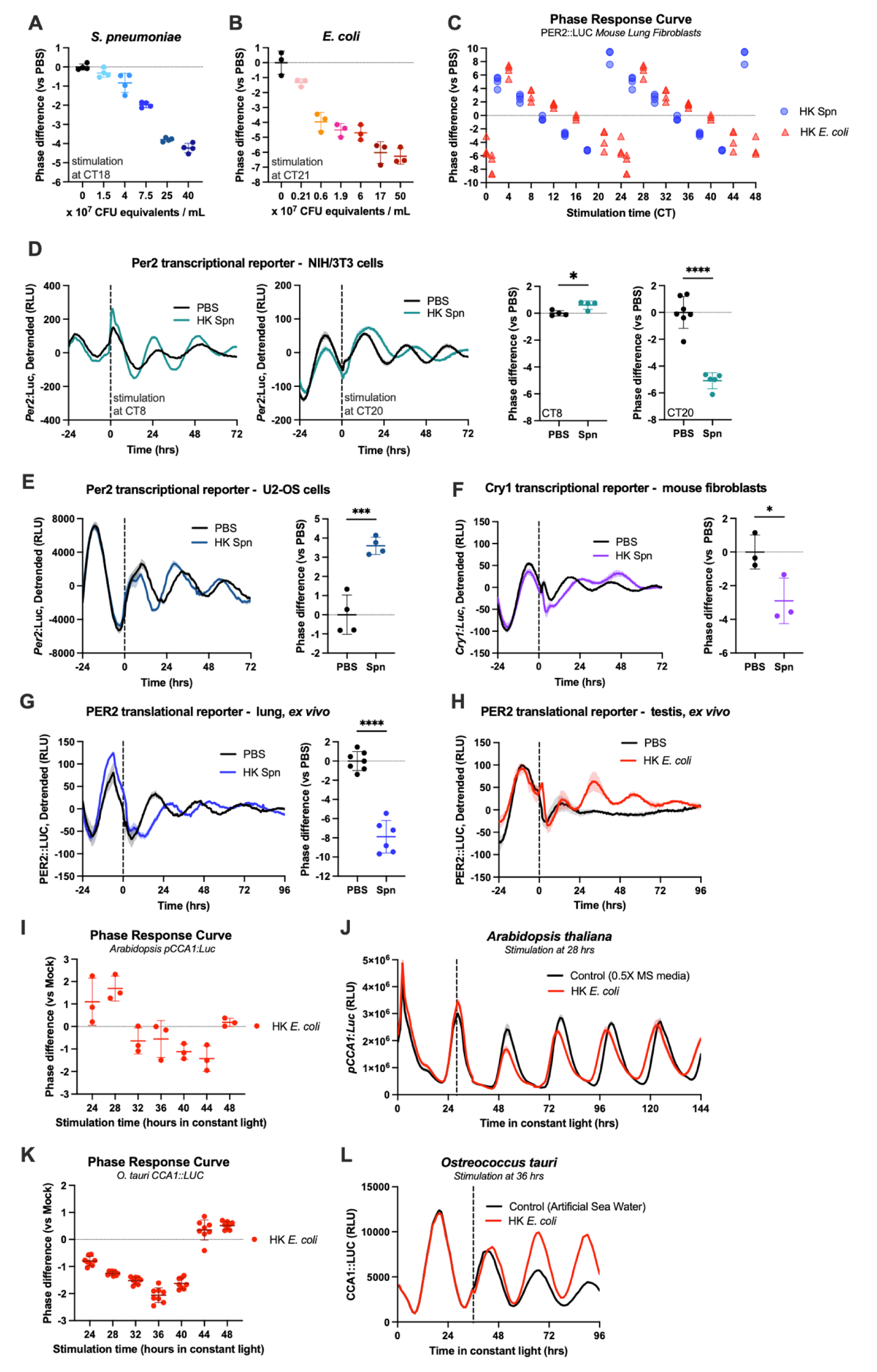
Bacterial signals reset circadian clocks across kingdoms. (**A-B**) P2L MLFs were stimulated with various concentrations of HK *Spn* (A) or *E. coli* (B) to generate a dose response curve, and the resulting phase difference compared to PBS control is graphed. Stimulation with 0 CFU equivalents / mL indicates the PBS control. (**C**) P2L MLFs were stimulated at various phases with either HK *Spn* or *E. coli* to generate a phase response curve (PRC). Data are double-plotted. A phase transition curve (PTC) and raw data are provided in Extended data Fig. 2. (**D**) *Per2:luc* transcriptional reporters in NIH/3T3 were stimulated with HK *Spn* or PBS at either CT8 or CT20 of *Per2* oscillation (dashed line). Each graph represents an individual experiment; samples are compared to the internal PBS control within that experiment. (**E-F**) U2-OS cells expressing a *Per2:luc* transcriptional reporter (E) and MLFs expressing a *Cry1:luc* transcriptional reporter were stimulated with HK *Spn* or PBS at time 0 (dashed line). (**G**) Organotypic lung slices from P2L mice were stimulated with HK *Spn* or PBS at time 0 (dashed line). (**H**) Organotypic testis slices from P2L mice were stimulated with HK *E. coli* or PBS at time 0 (dashed line). (**I-J**) *Arabidopsis thaliana* seedlings were stimulated with HK *E. coli* or media control at various CCA1 phases. The full PRC is shown in (I). The maximal change in phase occurred when bacteria were added at 28-hr in constant light (dashed line) (J). (**K-L**) *Ostreococcus tauri* were stimulated with HK *E. coli* or media control at various CCA1 phases. The full PRC is shown in (K). The maximal change in phase occurred when bacteria were added at 36-hr in constant light (dashed line) (L). Recording traces are shown as mean±S.E.; all other graphs are shown as mean±S.D. * = p<0.05; *** = p<0.001; **** = p<0.0001. *n* ≥ *3* for all groups, see Table S2 for all sample sizes and detailed statistical results.

We next asked whether this response extends across mammalian cell types. In NIH/3T3 murine embryonic fibroblasts expressing a *Per2:Luciferase* transcriptional reporter^39^, exposure to *Spn* resulted in a 1-hr phase advance when stimulation occurred at CT8 and a ∼5-hr phase delay when stimulation occurred at CT20). (Fig. 3D). As with the MLFs, the magnitude of the resultant phase shift was dependent on the phase at the time of stimulation. Human osteosarcoma cells (U2-OS) carrying a *Per2:luciferase*^39^ reporter (Fig. 3E) and mouse fibroblasts expressing a *Cry1:luciferase* reporter^34^ (Fig. 3F) also exhibit phase shifts, indicating that microbial cues can reset circadian timing across mammalian cell lineages and with different genetic reporters for the molecular clock. Finally, we exposed lung and testis tissue from P2L mice to HK bacteria *ex vivo* to determine if organotypic tissue slices also respond to microbial stimulation. Lung slices treated with *Spn* exhibited a robust ∼8-hr phase delay compared to PBS controls (Fig. 3G). While rhythmicity in PBS-treated testes damped too quickly to assess phase, *E. coli*-treated samples displayed restored oscillations, indicating a clear resetting cue upon microbial exposure (Fig 3H).

Circadian clocks across a wide range of organisms are often responsive by similar environmental signals^40^. To test whether microbe-induced circadian responses may be relevant outside of mammalian models, we examined the plant model *Arabidopsis thaliana* and the unicellular marine alga *Ostreococcus tauri*. Both species have transcriptional translational feedback loops (TTFL) involving proteins distinct from the mammalian clock. Using a transcriptional reporter of the promoter of *CIRCADIAN CLOCK ASSOCIATED 1* (*pCCA1:LUCIFERASE*) in *A. thaliana*, we monitored responses following stimulation with HK *E. coli*. Stimulation at various circadian phases revealed a maximal phase shift of ∼1.5-hrs (Fig. 3I-J). Using a translational fusion of CCA1 (CCA1::LUCIFERASE) in *O. tauri*, exposure to HK *E. coli* induced a maximal phase shift of ∼2-hrs (Fig. 3K-L). These findings suggest microbial exposure may serve as a relevant timekeeping cue in organisms beyond mammalian lineages.

### Circadian resetting is driven by a soluble factor distinct from canonical immune ligands

Our finding that diverse microbes elicit circadian phase shifts suggests a shared host signaling pathway may be involved. Pattern recognition receptors (PRR) such as Toll-Like Receptors and NOD-Like Receptors^41^ sense distinct microbial signals to activate common downstream inflammatory cascades^42^. We tested whether PRR agonists could replicate the observed phase shift; however, none of the tested ligands altered PER2 phase (Fig. 4A-C, Extended data Fig. 3A-E). Our P2L MLFs robustly produced IL-6 in response to these stimuli, confirming intact sensing capacity (Extended data Fig. 3F). We also tested purified *Spn* cell wall, a highly immunostimulatory complex containing peptidoglycan and wall-linked teichoic acids^43^, but this also failed to recapitulate the *Spn*-induced phase shift (Fig. 4D). Thus, neither activation of individual innate receptors nor inflammation is likely sufficient to drive cellular resetting. Finally, because the *Spn* cell wall remains intact following purification^44,45^, its lack of effect suggests that mechanosensory input is also insufficient to drive phase resetting.

**Figure 4.**
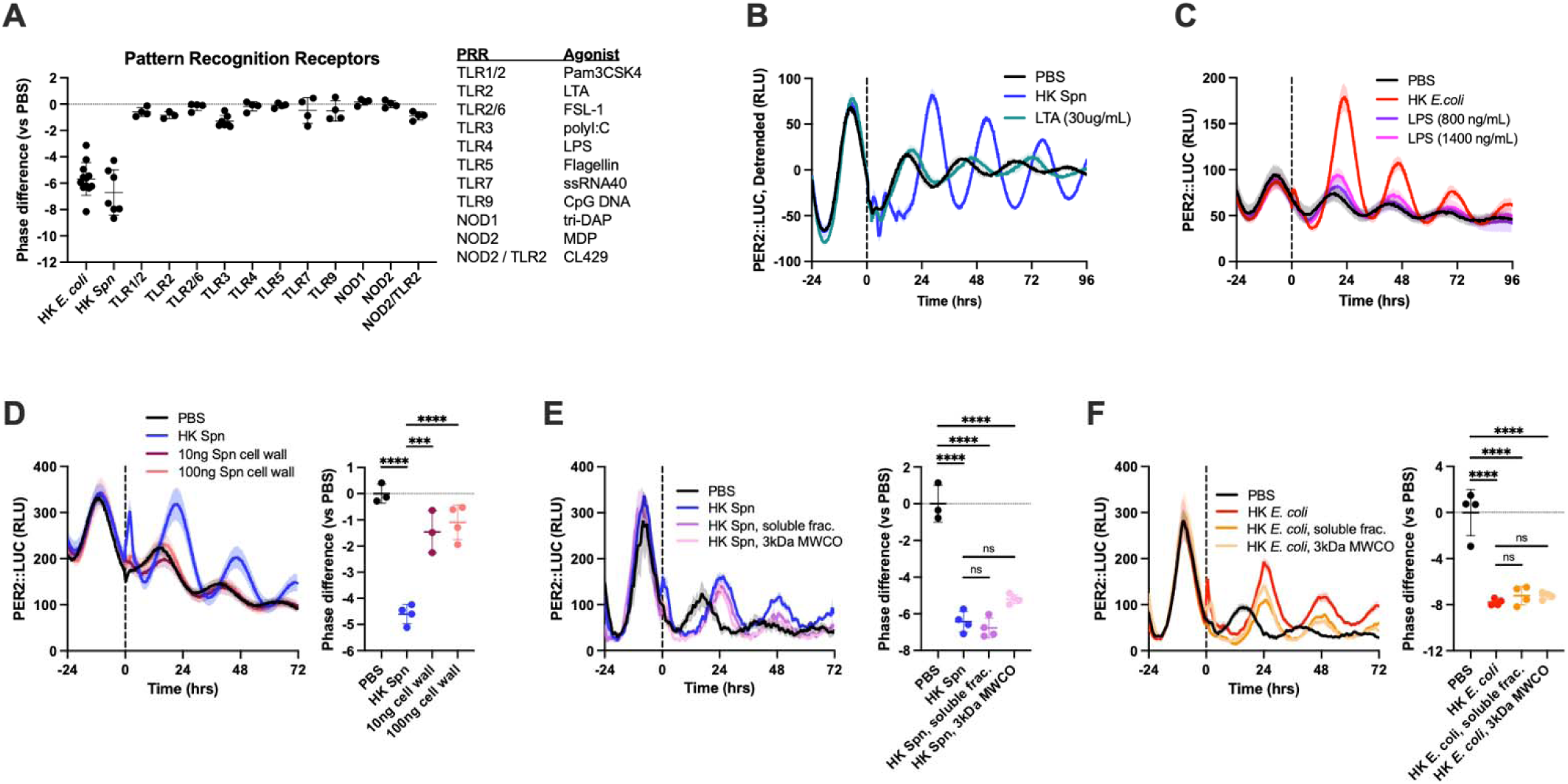
Circadian resetting is driven by a soluble factor distinct from canonical immune ligands. (**A-C**) P2L MLFs were stimulated with pattern recognition receptor (PRR) agonists across multiple independent experiments. Cross-experiment data of the resulting phase differences between each agonist, as compared to internal PBS control is compiled in (A). Raw data of experiments are shown for LTA (B) and LPS (C), with the remainder shown in Extended data Fig. 3. (**D**) P2L MLFs were stimulated with purified *Spn* cell wall and compared to PBS (negative control) and HK *Spn* (positive control). (**E-F**) HK *Spn* (E) or HK *E. coli* (F) were left as a crude mixture or pelleted by centrifugation and the soluble fraction (frac.) was collected. The soluble fraction was further passed through a 3kDa MWCO filter and the filtrate was collected. All samples were added to P2L MLFs at time 0. Recording traces are shown as mean±S.E.; all other graphs are shown as mean±S.D. *** = p<0.001; **** = p<0.0001; ns = not significant. *n* ≥ *3* for all groups, see Table S2 for all sample sizes and detailed statistical results.

To test whether mechanical stimulation is necessary for activity, we centrifuged HK *Spn* or *E. coli* bacteria and collected the soluble fraction. The soluble fraction induced similar PER2 phase shifts as HK bacteria, indicating that activity is associated with a component that is released upon lysis in both *Spn* and *E. coli* (Fig. 4E-F). We further passed these soluble fractions through a 3 kDa molecular weight cutoff (MWCO) filter and found that the filtrates retained full activity (Fig. 4E, F). Boiling releases cytoplasmic and some surface-associated molecules, though the effect varies as Gram-negative outer membrane components are more easily liberated than wall-anchored structures in Gram-positives. However, that activity is retained following boiling and filtration through a 3 kDa MWCO filter demonstrates that changes in PER2 phase can be driven by detection of heat-stable small molecule(s) produced by bacteria.

### p38 MAPK is a mediator of the bacteria-induced phase shift response

We next examined how bacterial exposure acts to induce PER2 by utilizing *Bmal1^−/−^* PER2::LUC cells, which lack transcriptional circadian cycles. Similar to previous experiments, bacterial exposure drove a rapid increase in PER2 that reached a maximum at several hours post-stimulation, suggesting that an intact circadian TTFL is not required for this response (Fig. 5A). To determine if nascent transcription is necessary for PER2 protein induction, we treated cells with [Z-amanitin, an inhibitor of RNA polymerase II and III, 30 minutes prior to *Spn* stimulation. PER2 levels still increased, indicating *de novo* RNA synthesis is not required for PER2 protein induction (Fig. 5B). Leptomycin B, which blocks nuclear mRNA export, showed similar results (Fig. 5C), suggesting reliance on transcripts already in the cytoplasm. In contrast, cycloheximide abolished acute PER2 induction, confirming translation is necessary for the observed induction (Fig. 5D). All three inhibitors caused PER2 rhythms to flatline within 12-hrs, consistent with the need for ongoing transcription and translation to sustain production of the short-lived PER2 protein^15^ (Fig. 5B-D). We do not exclude the possibility that transcriptional regulation of *Per2* is involved; however, our data suggests that there exists at least one contributing pathway which acts upon PER2 translation in a transcription-independent manner.

**Figure 5.**
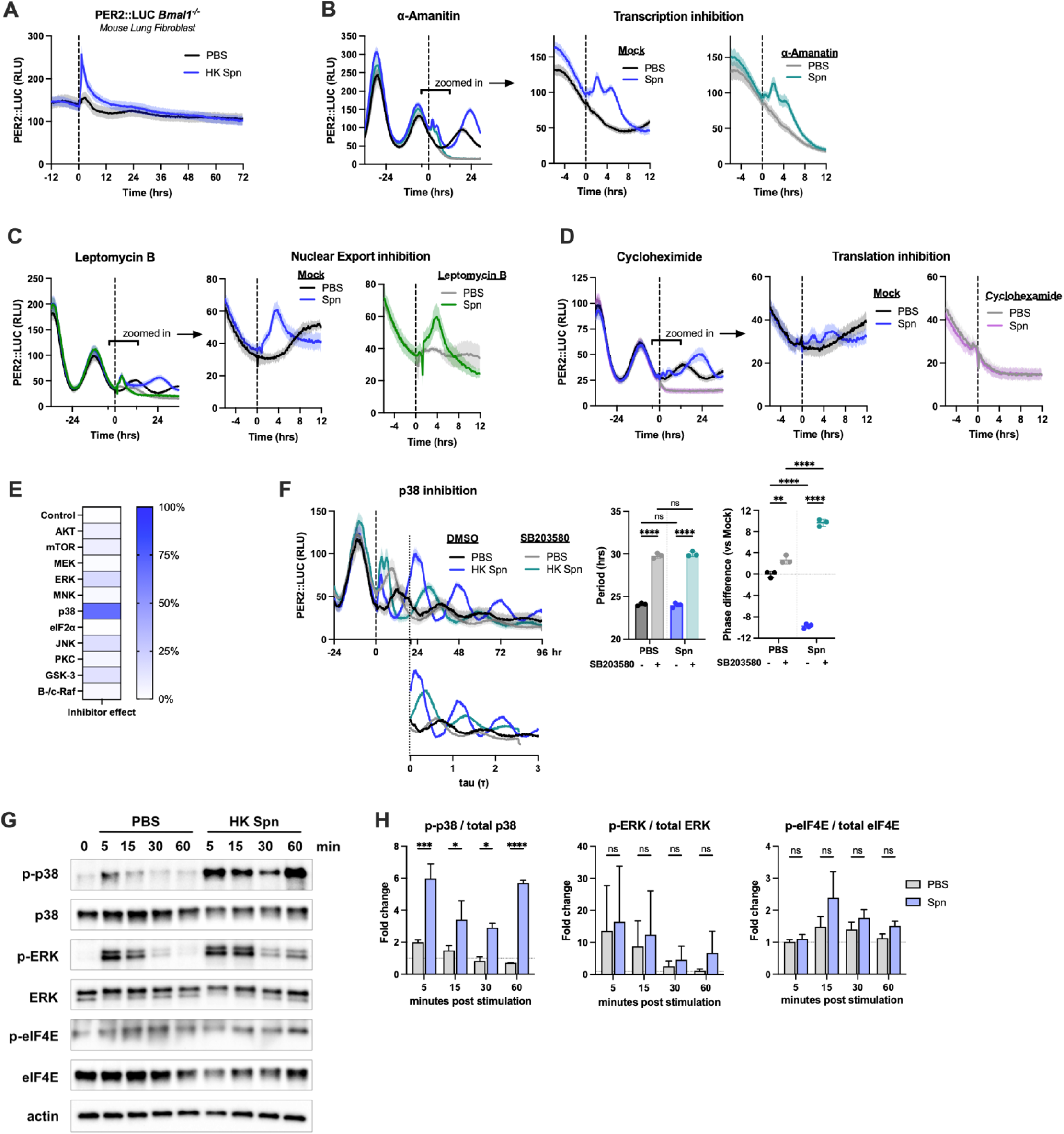
p38 MAPK is a modulator of bacteria-induced phase shift response. (**A**) *Bmal1^−/−^* PER2::LUC cells were stimulated with HK *Spn* or PBS at time 0. (**B-D**) P2L MLFs were pre-treated with vehicle (mock) or [Z-amanitin (B), leptomycin B (C), or cycloheximide (D) 30 minutes prior to stimulation with HK *Spn* at time 0 (dashed line). Zoom in of −6 to 12 hr is shown to visualize acute changes in luminescence. Relative Luminescence Units (RLU). (**E**) A heat map showing the relative ability of pharmacologic inhibitors to alter *Spn*-induced phase shifts. 0% represents no effect, while 100% represents a larger effect. (**F**) P2L MLF were treated with p38 MAPK inhibitor SB203580 30-min before stimulation with HK *Spn* or PBS at time 0 (dashed line, left). RLU is shown as unprocessed traces and, below, in modulo-*tau* format normalized based on period (τ). Modulo-*tau* graphs begin when the oscillations stabilize post-stimulation, indicated by the dotted line connecting the two graphs. Groups with a longer period (i.e. SB203580) end earlier in modulo-*tau* format as fewer circadian cycles were completed within 96 hr of recording. The effect of SB203580 on period and phase is shown. Phase difference is calculated in circadian hours compared to control samples (DMSO+PBS). (**G**) Synchronized P2L MLFs were stimulated with HK *Spn* or PBS at CT19 and protein samples harvested at 0, 5, 15, 30, or 60-min were analyzed by Western blot. Total and phosphorylated (activated) p38, ERK, and eIF4E are shown. Actin is a loading control. (**H**) Densitometry analysis of Western blots shown in (G) and Extended data Fig. 6A. The ratio of activated protein was calculated at each timepoint as [phosphorylated protein] / [total protein]. The fold change in activation was calculated by comparing each timepoint to the 0 minute sample on the same blot. Recording traces are shown as mean±S.E.; all other graphs are shown as mean±S.D. * = p<0.05; ** = p<0.01; *** = p<0.001; **** = p<0.0001; ns = not significant. *n = 2 for (H); n* ≥ *3* for all other groups. See Table S2 for all sample sizes and detailed statistical comparisons.

We next investigated whether known mechanisms of PER2 translational regulation may contribute to the response following microbial exposure. Previous studies demonstrate that miR-24, miR-29a, and miR-30a bind the 3’-UTR of *Per1* and *Per2* mRNA to suppress translation^46^. Downregulation of these miRNAs contribute to the resetting response to insulin, during which PER2 experiences a similar acute increase in abundance^34^. Following *Spn* stimulation, we found a modest decrease in the amount of miR-30a but not miR-24 or miR-29a (Extended data Fig. 4). As suppression of multiple miRNAs is typically required to enhance PER translation^34,46^, this limited change is unlikely to mediate the bacterial PER2 response.

Given the diversity of upstream signaling pathways capable of influencing the clock^47,48^, we next screened pharmacological inhibitors targeting diverse nodes of signal transduction. Because some compounds produced independent effects on phase or period, we established a prioritization metric to rank their impact on bacteria-induced resetting. For each inhibitor, we calculated the absolute value of the phase difference between *Spn*-treated MLFs in the presence or absence of the inhibitor. This approach captures both addition and reduction in phase as equivalent modulatory effects, and values are presented as a percent of the theoretical 12-hour maximum (Fig. 5E).

Interestingly, the mTOR inhibitor Torin-1 did not affect the phase shift response to *Spn* (Fig. 5E, Extended data Fig. 5A). mTOR is a central regulator of translation and has previously been shown to mediate circadian responses to temperature^49^, insulin^34^, and hypoxia^50^. As previously reported^51^, Torin-1 lengthens period (Extended data Fig. 5A *i*). To disentangle period effects from phase changes, we additionally visualized the PER2 traces in modulo-*tau* format, starting from when oscillations stabilize post-stimulation. Here, the data is normalized by the period of each group, such that the x-axis unit (τ) reflects 1 circadian cycle for all groups. This demonstrated that Torin-1 has no effect on the phase resulting from *Spn*-stimulation (Extended data Fig. 5A *ii*).

Of all the inhibitors tested, only SB203580, a p38 MAPK inhibitor, significantly altered the response to *Spn* (Fig. 5F, Extended data Fig. 5B). Like Torin-1, SB203580 lengthened the period in both PBS- and *Spn*-treated cells. However, in contrast to Torin-1, the phase shift induced by *Spn* was markedly altered in the presence of SB203580 (Fig. 5F, Extended data Fig. 5A-B). Additional studies demonstrated that BIRB796, a structurally distinct molecule that inhibits p38 MAPK, but not an inhibitor of ERK, also leads to an altered *Spn*-induced phase shift (Extended data Fig. 5C-D). Given that phase is cyclical, it is not intuitive whether the final phase results from inhibition or augmentation of the *Spn* response as compared to PBS controls. However, when the phase difference between (*Spn* + p38 inhibitor) and (*Spn* + mock control) is calculated, all p38 inhibitor experiments show an additional phase delay, demonstrating a consistent effect of p38 inhibition (Extended data Fig. 5E). In contrast, inhibition of mTOR or ERK has no significant effect.

Our inhibitor experiments suggest that *Spn* may be engaging the host p38 MAPK pathway to mediate the phase shift effect. We thus tested the effect of p38 MAPK activation on PER2 phase. Anisomycin, a p38 MAPK activator, induced a phase delay similar to addition of HK *Spn*, suggesting that p38 MAPK activation alone can replicate the phase shift effect (Extended data Fig. 5F). Although evidence for p38 MAPK in mammalian circadian control is limited in the literature, related MAPK signaling pathways have been implicated in clock resetting. Specifically, ERK participates in light-induced phase shifts through the ERK–MNK–eIF4E axis, a translational control mechanism that culminates in phosphorylation of the cap-binding protein eukaryotic Initiation Factor 4E (eIF4E)^52^. Phosphorylation of eIF4E (p-eIF4E) is further linked to circadian oscillations in locomotor activities^52^ and cognitive functions^53^, implicating it as a potential mediator of diverse clock outputs. While our inhibitor screen suggested ERK is not involved in the response to bacteria (Extended data Fig. 5D), p38 can also activate MNK to phosphorylate eIF4E^54,55^. Thus, we investigated if the phase shift to bacterial exposure may be mediated by a p38–MNK–eIF4E axis.

We asked whether *Spn* engages this axis by examining phosphorylation status following microbial stimulation. Phosphorylation of p38 is induced by *Spn* within 5 minutes and remains elevated 60 minutes later (p-p38, Fig. 5G-H, Extended data Fig. 6A). In contrast, ERK phosphorylation transiently increases in both *Spn*- and PBS-treated cells before returning to baseline, consistent with a generalized handling response rather than a microbe-specific signal. We next tested whether p-p38 might act through a canonical translational pathway involving MNK and eIF4E, as was described for light^52^. However, we did not detect changes in p-eIF4E following *Spn* exposure, suggesting that microbial stimulation activates a distinct translational mechanism that bypasses this axis. These findings, combined with our pharmacological studies, point to p38 MAPK as a modulator of the bacterial resetting response, operating outside established translational axes linked to circadian control. More broadly, these findings suggest that bacterial exposure can engage a noncanonical signaling mechanism to rapidly influence circadian timing, distinct from previously characterized clock-resetting pathways.

### Repeated bacterial exposure continuously modifies circadian timing

Although microbes are ubiquitous in the environment, their levels and community composition is highly dynamic. Microbial loads in the atmosphere follow a daily rhythm, with a 10-100 fold greater abundance reported at night as compared to the day, due to reduced atmospheric mixing^13^. The gut microbiome shows a comparable rhythmicity, with up to 15% of taxa oscillating daily in concert with host feeding rhythms^14^. This raised the question of whether continuous presence of microbes, combined with modest changes in microbial abundance are sufficient to alter the behavior of a circadian oscillator. To address this, we constructed a simple model of a hypothetical oscillator (Extended data Fig. 7A). In this framework, an activator protein A represents the positive arm of the TTFL, while a repressive protein B represents the negative arm.

As shown in this study, bacterial exposure is capable of inducing PER2 (part of the negative arm of the mammalian clock) in a *Bmal1*-independent manner (part of the positive arm). We therefore chose to model bacterial exposure such that an increase in bacterial signal directly induces repressor B independently of enhancer A. *In silico* modeling of a single bacterial exposure was able to produce a 4-hr phase shift (Extended data Fig. 7B), which is similar to the phase shift observed when bacteria are applied at a similar circadian phase (Fig. 3C, Extended data Fig. 2C-D). We next adjusted bacterial concentrations so that the signal was always present but oscillated with a 24-hr rhythm to simulate bacterial fluctuations in the environment. All simulations modeled a ten-fold increase over a minimum, baseline amount (1, 5, or 10 bacteria) which leads to differing amplitudes of bacterial oscillation (amplitude of 10, 50 or 100). We found that in all simulations, the *in silico* oscillator adjusted its phase over time to synchronize with the bacterial fluctuations (Extended data Fig. 7C), though changes in phase occurred at a greater rate when higher amounts of bacteria were simulated (Extended data Fig. 7D).

Previous studies examining the properties of *in vitro* circadian recording have shown that individual cells exhibit stable circadian oscillations, yet across a community of cells, stochastic fluctuations cause individual cells to deviate over time from the overall average, leading to an observed loss of amplitude over time^56–58^. Therefore, to model this community dynamic, we simulated random fluctuations in the circadian oscillator model, then averaged the measurements of 3000 independently simulated oscillators. When this community model was simulated with bacterial exposure at repeated 24-hr intervals, the model predicted that the community remains synchronized and resists amplitude loss (Extended data Fig. 7E).

We tested this model experimentally by evaluating whether repeated exposure to HK *Spn* every 24-hrs is sufficient to maintain synchrony in P2L MLF. As shown in Figure 6, repeated stimulations with HK *Spn* allowed P2L MLFs to maintain their amplitude for the duration of stimulation, after which damping occurred in a manner consistent with desynchronization. PBS (bacteria-free; handling control) had no effect at any timepoint, and amplitude in these cells damped with kinetics similar to untreated cells (grey vs light blue traces in Fig. 6B-D). In contrast, additional stimulation with HK *Spn* resulted in maintenance of amplitude (dark blue traces compared to grey or light blue in Fig 6A-D). As we did not remove HK *Spn* during the experiment, the observation that MLFs were able to respond to subsequent stimulations suggests that MLFs sense changes in bacterial abundance rather than merely the presence or absence of bacteria. This is consistent with the prediction that daily fluctuations in bacterial abundance are sufficient to influence mammalian circadian clocks. As microbial exposure is ubiquitous on Earth and as bacteria are known to oscillate in response to well-established entraining cues such as light, temperature, and feeding rhythms, we propose that diverse organisms may use bacterial detection as a mechanism to reinforce circadian synchrony with external time.

**Figure 6.**
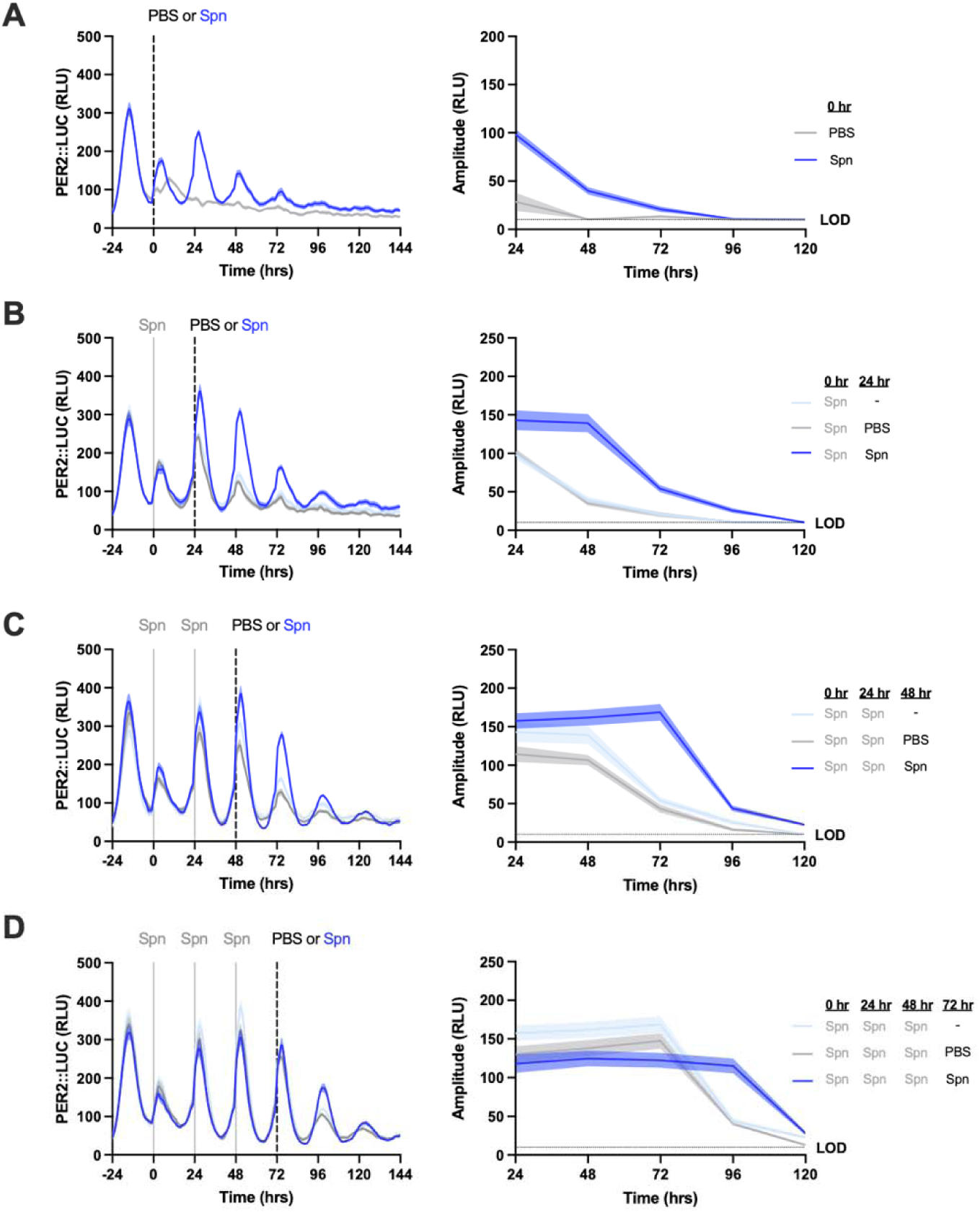
Repeated exposure to microbes maintains circadian amplitude. P2L MLF were stimulated with HK *Spn* or PBS at times 0-, 24-, 48-, and 72-hrs. Upon receiving PBS, P2L MLFs do not receive stimulation in subsequent stimulations. Recording traces from the same experiment are shown. The amplitude over time of the graphed traces is shown. (**A**) Recording traces of P2L MLF receiving HK *Spn* or PBS at time 0 (dashed line). (**B**) Recording traces of P2L MLF receiving HK *Spn* or PBS at time 24 (dashed line). The recording trace of P2L MLF receiving HK *Spn* at time 0 with no subsequent stimulation is shown as a no-handling control. (**C**) Recording traces of P2L MLF receiving HK *Spn* or PBS at time 48 (dashed line). The recording trace of P2L MLF receiving HK *Spn* at time 24 with no subsequent stimulation is shown as a no-handling control. (**D**) Recording traces of P2L MLF receiving HK *Spn* or PBS at time 72 (dashed line). The recording trace of P2L MLF receiving HK *Spn* at time 48 with no subsequent stimulation is shown as a no-handling control. All graphs are shown as mean±S.E. *n = 6* for all groups. See Table S2 for all sample sizes and detailed statistical comparisons.

## Discussion

Our findings reveal that microbial exposure can serve as a cue capable of resetting the mammalian clock. Microbial exposure thus emerges as a novel circadian clock input, joining well-characterized cues such as light^16^, temperature^17^, and feeding^18^ in shaping biological timing. The observation that similar effects occur in plants and algae suggests that sensitivity to microbial cues may be a general feature of biological timekeeping across kingdoms.

Across ecosystems, microbial communities fluctuate predictably in response to environmental changes, offering a rich and ancient source of temporal information^7–12^. Microbial cues may therefore act as circadian amplifiers – reinforcing or substituting for classical zeitgebers, especially in environments where cues like light or temperature are weak or inaccessible (e.g., soil, internal tissues). Notably, germ-free mice exhibit disrupted circadian oscillations despite standard light–dark housing^21^. While the extensive metabolic and immune defects present in germ-free mice likely contribute to perturbations in overall circadian rhythmicity *in vivo*, the data presented here demonstrate microbial signals *in vivo* may also play a direct role in clock maintenance.

In principle, circadian responsiveness to microbes could also confer local advantages. During infection, bacteria can reach high densities within host tissues, resulting in rapid exposure to microbial signals. The ability of host cells to adjust cellular phase upon detection of microbial signals could reflect an adaptive alignment that optimizes immune or stress responses without disturbing systemic rhythms. Given that pathogens engage host tissues through diverse mechanisms, resolving this question will require distinct analyses appropriate for each infection model. Notably, during *Spn* infection, high bacterial density triggers regulated bacterial lysis through at least two genetically separable pathways, LytA-mediated autolysis^59,60^ and Blp-dependent bacteriocin activity^61,62^, which together cause the release of intracellular bacterial components into the local environment. An important open question is therefore whether microbial-derived phase shifts occur during natural infections, and if so, how this alters disease outcome.

While these hypotheses are tempting to speculate over, such ideas must also be qualified by the current lack of understanding in the field of circadian host-microbe interactions^63^. For instance, although we have not observed differences in the bacteria-induced phase shift response across our tested mammalian models, it is possible that specific organs or cell types may respond to microbial exposure differently than we have observed *in vitro*. Additionally, recent research has revealed that some species of non-photosynthetic bacteria – such as *Bacillus subtilis*^64^, *Klebsiella aerogenes*^65,66^, and *Acinetobacter baumanii*^67^ – contain circadian clocks. While circadian clocks have not been reported in our representative bacterial species (*Spn* and *E. coli*), it is possible that one may be revealed in future studies. If an intrinsic circadian oscillator is discovered in these organisms, an exciting future direction would be to determine if it influences the microbe-induced phase shift. Understanding the biological function and evolutionary benefit of the observed phase shift effect will therefore require collective effort to explore this emerging interface between microbial biology and circadian timing.

Our studies point to a soluble bacterial factor as the effector of circadian resetting. While this activity is not mimicked by canonical PRR ligands, our data do not exclude the possibility of a combinatorial signal. Given the vast repertoire of microbial products and potential for synergistic interactions, identifying the precise signal(s) will be an important avenue for future investigation. Functionally, microbial stimulation triggers acute PER2 induction, but whether this drives the observed phase shifts or reflects a parallel consequence of upstream signaling remains unclear. Notably, while acute changes in transcriptional *Per2* reporters can be observed, we show that protein PER2 induction does not require *Bmal1*, transcription, or canonical translational control through mTOR or eIF4E. Given the complexity of MAPK signaling and the wide array of downstream targets^68^, it remains unresolved whether p38 potentiates, constrains, or gates the phase-resetting response^69^. These data define mechanistic constraints on the response but do not resolve the upstream sensing mechanism or the direct p38 targets responsible for PER2 induction. Unbiased approaches such as proteomic or phosphoproteomic profiling will be a valuable future direction to define the broader signaling landscape linking bacterial exposure to regulation of the molecular clock. Microbial exposure is well known to influence a wide range of host cellular processes beyond circadian regulation; however, how these pathways intersect with the clock-resetting response described here remains to be defined.

Together, these findings position microbial signals as a novel class of circadian input and uncover a previously unrecognized axis of host–microbe communication. That this signaling appears independent of classical sensing or clock-resetting pathways suggests the existence of a broader and more diverse landscape of circadian entrainment than previously appreciated. This interaction between bacterial signals and host rhythmicity opens exciting questions about its origins and functional relevance. We hope these findings lay the foundation for future studies into how microbial cues shape circadian timing across commensal, environmental, and disease contexts.

## Supporting information

Table S1 (Bacterial growth conditions)

## Acknowledgments

We would like to thank the following individuals for providing resources: Dr. Jason Rosch (*Spn* strains A66.1, BHN97, TIGR4 Δ*cps*), Dr. Elaine Tuomanen (*Spn* cell wall extract), Dr. Samantha Bell (heat killed *Listeria monocytogenes* and *Mycobacterium smegmatis)*, and Dr. David Welsh (P2L mouse lungs). We would also like to thank the staff of MRC ARES for their assistance with animal work.

## Funding

NIH/NIGMS grant R35GM147509 (JMK)

Startup funds provided by UC Santa Cruz (JMK)

Pew Scholar in the Biomedical Sciences as part of the Pew Charitable Trusts (JMK)

Hellman Fellows Program (JMK)

Alfred P. Sloan Foundation (JMK)

NIH/NIGMS grant R35GM141849 (CLP)

Howard Hughes Medical Institute (CLP)

BBSRC (UK) BB/Z516636/1 (AND)

Institute Strategic Programme BRiC BB/X01102X/1 (AND)

European Research Council Synergy award 10116698 “MicroClock” (AND)

Boehringer Ingelheim Fonds travel grant (PC)

Wellcome Trust Discovery Award 225212/Z/22/Z GvO)

## Authors contributions

Conceptualization: DM, PC, JMK

Investigation: DM, DNL, PC, EB, OPF, GvO, JMK

Methodology: DM, PC, JMK

Validation: DM

Formal analysis: DM, PC, EB, GvO, JMK

Resources: CLP, AND, JD, GvO, JSO, JMK

Data curation: DM, TNL, PC, JMK

Writing – original draft: PC, DM, JMK

Writing – review & editing: DM, AND, JMK

Visualization: DM, JMK

Supervision: CLP, AND, GvO, JMK

Funding acquisition: CLP, AND, GvO, JSO, JMK

Project administration: JMK

## Competing interests

Authors declare that they have no competing interests.

## Data and materials availability

All data are available in the main text or the supplementary materials.

## Supplementary Materials

### Materials and Methods

#### Bacterial strains and preparations

Detailed information on bacterial strains, phenotypes, media type and culture conditions can be found in **Table S1**. Briefly, most bacterial strains were inoculated into overnight liquid culture, then subcultured and grown to mid log (OD_600_ 0.4-0.5) with the exception of *Streptococcus pneumoniae*, which was always maintained in log phase (to prevent autolysis that occurs in stationary phase), and *E. coli* DH5α which exclusively used stationary phase cultures. *Streptococcal* species (*S. pneumoniae* and *S. agalactiae)* were grown in liquid media without agitation. All other bacteria were grown under shaking conditions when in liquid culture. Bacterial colony forming units (CFU) were enumerated by 10-fold serial dilution and counted following overnight growth on solid agar media. For experiments involving inactivated bacteria, CFU was calculated prior to inactivation and is subsequently referred to as “CFU equivalents”.

**Heat Killed (HK)** bacteria were grown as described and a sample was collected for CFU enumeration. Cultures were then pelleted by centrifugation and resuspended in minimal PBS (for mammalian cells), half-strength Murashige and Skoog media (for *A. thaliana*), or salt water (for *O. tauri*), typically resulting in at least 100-fold concentration (resuspend 50mL bacterial culture in < 500uL). Bacteria were heat killed by boiling in a heat block at 100C for 30-60 min or until plating revealed no viable bacteria. As *B. subtilis* produces heat-resistant endospores, these samples were additionally autoclaved after boiling. For fractionation experiments, HK *Spn* or *E. coli* were centrifuged at max speed (21,000 x g) to separate insoluble pellet from the soluble fraction. The soluble fraction was then passed through a 3kDa MWCO filter (Millipore Sigma UFC5003) according to manufacturer’s directions. All samples were stored at −20°C until use.

**Formalin Fixed (FF)** bacteria underwent identical enumeration and concentration steps as HK bacteria. After resuspension in PBS, bacterial pellets were diluted in 10% Neutral Buffered Formalin and incubated at room temperature for 15 minutes. Formalin was then quenched and removed by thorough washing with 3M Tris-HCl pH 8 (a minimum of three 15-minute washes was necessary to prevent formalin-mediated toxicity in downstream steps), and bacterial pellets were resuspended in minimal PBS. Neutralization of formalin was confirmed by adding formalin-inactivated bacteria to confluent P2L cells at a final concentration of 5% (v/v) and checking for the lack of morphological signs of cell stress over a period of 48 hours.

#### Eukaryotic cell culture

Mouse lung fibroblasts (MLF) were isolated as described previously^34^ from PERIOD2::LUCIFERASE^22^ (P2L) mice. The NIH/3T3 murine embryonic fibroblast, U2-OS human osteosarcoma, and *Cry1:luciferase* mouse fibroblast cell lines were produced previously^34,39^. All cells were grown in DMEM (Gibco 10569044) supplemented with GlutaMAX (Gibco 35050061) and 10% (v/v) FBS (Hyclone FetalClone III, ThermoFisher) and maintained with regular media changes.

MLFs were seeded and allowed to reach full confluency before use in any assay. As these cells exhibit strong contact-inhibition, variations in initial seeding density and time to confluency did not have any noticeable effects on downstream experiments. NIH/3T3 cells were seeded at 10% confluency used within 2 weeks of seeding. U2-OS cells were seeded at 50% confluency and used within 3 days of seeding.

#### Bioluminescence recordings of mammalian samples

Bioluminescence recordings of mammalian cells and organotypic slices occurred in either a LumiCycle 32 (Actimetrics), Tecan Spark (Tecan), or ALLIGATOR (Cairn Research). All cell lines were synchronized by temperature cycles of 12 hours at 37°C followed by 12 hours at 32°C from seeding. Immediately before recording, cells were refreshed into the appropriate recording media. To maintain the entrainment with the prior temperature cycling regimen, media changes occurred between 6 to 10 hours after a 32°C-37°C temperature shift.

#### Recording media (CO_2_)

DMEM (Gibco 31053036) supplemented with GlutaMAX (Gibco), sodium pyruvate (Gibco), 10% FBS, and 1 mM D-luciferin (potassium salt, CAS# 115144-35-9, Biosynth International FL08608)

#### Recording media (no-CO_2_)

DMEM powder without sodium bicarbonate (Sigma D2902-1L) supplemented with 4 mg/mL additional glucose, 0.35 mg/mL sodium bicarbonate, 0.02 M MOPS, 100 ug/mL penicillin/streptomycin (Millipore Sigma P4333-100ML), 2% (v/v) B27 or NS21, and 1 mM D-luciferin (potassium salt, CAS# 115144-35-9, Biosynth International FL08608), pH 7.4, 325-340 mOsm

During stimulation, mammalian cells and tissues were removed from the recording device on a veterinary heating pad (Deltaphase Isothermal Pad, Braintree Scientific) and kept at a constant 37°C for the duration of the treatment before returning to the recording device. Handling times were kept as short as reasonably possible, with handling times of larger experiments spanning up to 5 minutes. The volume of the stimulants added was at or less than 5% of the volume of the cell culture media to minimize temperature disruptions. In experiments where a wash was performed, cells were placed in recording media without serum. During the wash, media was removed from the cell culture, the cells were washed with warm PBS, and fresh recording media was added to the cell culture.

#### LumiCycle

Cells were seeded in tissue culture-treated 35 mm dishes and maintained as described above. Immediately before recording, the media was refreshed into recording media (no CO_2_). Dishes were sealed with glass cover slips and vacuum grease (Dow Corning 1597418) and placed into the LumiCycle at constant 37°C.

#### Tecan Spark

Cells were seeded in tissue culture-treated 24- or 48-well plates. Immediately before recording, the media was refreshed into recording media (CO_2_), and wells at the plate edge were filled with PBS to serve as a humidity buffer. Plates were placed in the Tecan Spark at constant 37°C, 5% CO_2_, and bioluminescent activity was recorded with a 40 second per well exposure time and recording photon counts (not normalized to time).

#### ALLIGATOR

Cells were seeded in tissue culture-treated 24-, 48-, or 96-well plates or 35 mm dishes. Immediately before recording, cells were placed in appropriate recording media depending on whether or not CO_2_ was provided during the experiment. Bioluminescent activity was then recorded using the electron multiplying charge-coupled device (EM-CCD) on the ALLIGATOR. ALLIGATOR recordings were processed with the Cosmic Noise Reducer FIJI plugin^70^ and quantified with FIJI^71^.

#### Organotypic slices

All animal experiments were licensed under 1986 Home Office Animal Procedures Act (UK) and carried out in accordance with local animal welfare committee guidelines. Adult PER2::LUC^22^ mice were euthanized by cervical dislocation and confirmed by exsanguination. Slices were sectioned to 300 μm thickness in ice-cold phosphate-buffered saline (PBS) using a McIlwain Tissue Chopper (Stoelting) and placed in 1.2 mL DMEM (Gibco, D5921), supplemented with 1% (v/v) GlutaMAX (Gibco), 2% (v/v) NS21, penicillin/streptomycin, and 1 mM D-luciferin. Tissue slices were recorded in an ALLIGATOR (Cairn Research) at constant 37°C immediately upon sectioning.

#### *Arabidopsis thaliana* propagation and recording

For experiments with the model plant *Arabidopsis*, seedlings of *Arabidopsis thaliana* (Col-0 background, harboring the transcriptional reporter of CIRCADIAN CLOCK ASSOCIATED 1 (*pCCA1:LUCIFERASE*)) were cultivated in dense clusters, as described previously, using sterile tissue culture^72^. Surface-sterilized seeds were stratified for 3 days at 4°C in darkness, then cultivated under cycles of 12-hr light and 12-hr dark, 19°C, for 10 days, under 80 μmol m^−2^ s^−1^ of white light.

On day 10, 5 mM D-luciferin (potassium salt; Biosynth) was applied to seedlings, which were transferred on day 11 to a Photek EM-CCD bioluminescence imaging system under 34 μmol m^−2^ s^−1^ red and blue light supplied with custom LED panels. Heat-inactivated *E. coli* stocks were diluted in half-strength Murashige & Skoog media (plant growth media; Duchefa) and applied at 4-hr intervals, over 24-hrs, to separate batches of seedlings, with additional seedlings receiving a mock treatment as a phase reference. Subsequently, *pCCA1:LUCIFERASE* bioluminescence was recorded for 7 days under constant light conditions^72^.

#### *Ostreococcus tauri* culture and recording

For experiments with Ostreococcus tauri, a transgenic line expressing the CCA1 gene, fused to firefly luciferase, from its native promoter was used^73^ (CCA1::LUCIFERASE). Cells were cultured in artificial sea water as reported previously^74^ and subjected to cycles of 12-hr light and 12-hr dark under 15-20 μmol m^−2^ s^−1^ of blue light (Lee Filters 183 “Moonlight Blue”). 6 day old cultures were transferred to 384 well plates (Greiner Bio-One Lumitrac). After 3 days in entrainment, 1 mM D-luciferin (potassium salt; Biosynth) was added, and imaging was started under constant light conditions at dawn on the 4th day using a Tristar2 plate reader (Berthold) as previously reported^75^. Heat-inactivated *E. coli* stocks and controls were applied at 4-hr intervals to replicate wells, and bioluminescence was recorded every hour.

#### Analysis of bioluminescence traces

In experiments where the baseline of bioluminescent traces was linear, traces from mammalian cells were fitted with damped cosine waves in Prism 10 (GraphPad) using the following equation:

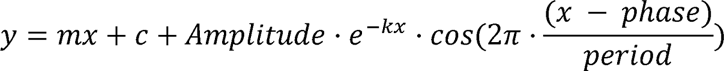

where y is the luminescence, x the time, amplitude is the height of the peak of the waveform above the trend line, k is the exponential decay rate, phase is the horizontal shift of the wave, and the period is the time taken for a complete cycle to occur.

In experiments where the baseline was non-linear, fitting with this equation was inaccurate, thus the bioluminescent traces were detrended until a linear baseline was achieved. A 24-hour moving average of each sample was calculated to estimate the baseline of the sample, and the baseline was further refined by fitting the 24-hour moving average to an exponential decay, linear, quadratic, or cubic equation in Prism 10. This baseline was then subtracted from the measured luminescence.

We observed that the bioluminescence traces of P2L MLFs stabilized over the course of approximately 16 hours post-stimulation and those of NIH/3T3 and U2-OS stabilized over the course of approximately 24 hours. Therefore, the phase following bacterial stimulation was calculated when the traces stabilized, which typically occurred between 12 to 24 hours post-stimulation for P2L MLFs and 24 hours post-stimulation for NIH/3T3 and U2-OS cells. Statistical analyses of quantified bioluminescent traces were performed using Prism 10.

With *A. thaliana*, we observed that after treatment with heat-inactivated *E. coli*, a transient change in period occurred over a duration of about 24 h, until a new stable state was reached. Therefore, to construct PRCs, the phase following HK *E. coli* treatment was estimated between 72 hours and 144 hours of free run using FFT-NLLS analysis in BioDare2^76^ (biodare2.ed.ac.uk). With *O. tauri*, phase was calculated from treatment time to 96 hours of free run.

The fitted phase was used to calculate the phase at the beginning of the measurement window, which was used for phase and phase difference in all experiments. Phase is reported in circadian time (CT), where an oscillation trough is defined as CT0, 24 circadian hours is defined as the period of the oscillation, and CT3 corresponds to 3 circadian hours after CT0. Phase difference is reported as the difference in phase between the test stimulation (e.g. bacteria) and control (e.g. PBS) groups in circadian hours, such that a negative phase difference represents a phase delay relative to the control and a positive phase difference represents a phase advance relative to the control.

#### Pharmacological treatment

For experiments using inhibitors, the recording media were modified to replace FBS with 2% NS21 or B27 and recorded as otherwise described above. Inhibitors were prepared in DMSO such that the final concentration of DMSO in the cell culture media was at or less than 0.1% (v/v), and the inhibitors were added to the cell culture 30 minutes before stimulation. Final concentrations of the inhibitors are as follows: PKC (Sotrastaurin, 500 nM), AKT (MK 2206, 10 μM), mTOR (Torin-1, 1 μM), MEK (U0126, 10 μM), JNK (SP 600125, 3 μM), ERK (SCH772984, 100 nM), p38 MAPK (SB203580, 10 μM), MNK1 (CGP 57380, 10 μM), eIF2[Z (ISRIB, 200 nM), B-/c-Raf (Sorafenib, 3 μM), GSK-3 (CHIR99021, 3 μM). For experiments with BIRB796 (second p38 inhibitor), BIRB796 was added to a final concentration of 10 μM. As with the stimulation procedure, cells were kept on a veterinary heating pad during the inhibitor treatment. For experiments with anisomycin (p38 activator), anisomycin was added to a final concentration of 0.1 μM at the time of stimulation.

#### Pattern Recognition Receptor Agonists

Bacterial cell wall was donated by Elaine Tuomanen and Jason Rosch and was extracted from *Spn* via an established protocol^44,45^. The remaining ligands were purchased from commercial vendors. TLR1/2, TLR3, TLR4, TLR5, TLR2/6, TLR7, and TLR9 agonists (Mouse TLR1-9 Agonist kit, Invivogen tlrl-kit1mw), NOD1 agonist (Invivogen tlrl-tdap), NOD2 agonist (Invivogen tlrl-mdp), and NOD2/TLR2 agonist (Invivogen tlrl-c429) were purchased from Invivogen. TLR2 agonist was obtained from Millipore Sigma (L3265-5MG).

#### qRT-PCR for mRNA

P2L MLFs were grown to confluence in 35 mm dishes and synchronized via temperature cycling as described above. A paired set of dishes were handled similarly and recorded for bioluminescent traces as described above. Before recording, the media for all dishes was changed into recording media (no CO_2_) without FBS or NS21. After treatment, cells were harvested and RNA was extracted using the RNeasy Mini Kit (Qiagen) according to manufacturer’s instructions. cDNA was generated from these samples using the iScript cDNA synthesis kit (Bio-Rad) using a total RNA concentration of 100 ng per sample. Samples were then kept on ice while they were made up to a volume of 100 μL with nuclease-free water. Small quantities of all samples were pooled and subsequently serially diluted to make standards. Samples were diluted tenfold to ensure their values fell within the standard curve. qPCR was carried out using a QuantStudio 3 machine (Applied Biosystems) and PerfeCTa SYBR Green FastMix (Quantabio) as per manufacturer’s protocols. Primer sequences used are as follows: *Per2* forward 5’-CCTACAGCATGGAGCAGGTTGA-3’, *Per2* reverse 5’-TTCCCAGAAACCAGGGACACA-3’, *Rn18s* forward 5’-CGCCGCTAGAGGTGAAATTC-3’, *Rn18s* reverse 5’-TTGGCAAATGCTTTCGCTC-3’. All pairs of primers were annealed at 58°C. Data was then analyzed using the QuantStudio 3 software (Applied Biosystems). All values were normalized against RN18S.

#### qRT-PCR for miRNA

P2L MLFs were harvested and miRNA isolated using an miRNeasy Micro Kit (Qiagen) as per the manufacturer’s protocol. On-column DNase digestion was not performed. cDNA was generated using the miScript II RT Kit (Qiagen). Briefly, a total reaction volume of 20 μL consisted of 4 μL HiSpec Buffer, 2 μL miScript Nucleics Mix, 2 μL miScript Reverse Transcriptase Mix, and 500 ng RNA, made up to volume with nuclease-free water. Samples were then incubated at 37°C for 60 mins, followed by 5 min at 95°C. Samples were kept on ice while they were made up to a volume of 100 μL with nuclease-free water. Pooled standards and further dilution of samples was performed as for qPCR for mRNA. qPCR was performed using the miRCURY LNA SYBR Green PCR Kit (Qiagen). Briefly, wells consisted of 5 μL miRCURY SYBR Green PCR Master Mix, 1 μL PCR primer mix, and 1 ng cDNA, made up to 10 μL with nuclease-free water. The plate was then sealed, vortexed and centrifuged briefly before being placed in the QuantStudio 3 Real-Time PCR machine (Applied Biosystems). Wells were then subjected to 15 min at 95°C, followed by 40 cycles of 15 seconds at 94°C, 30 seconds at 55°C, then 30 seconds at 70°C. Results were analyzed using the QuantStudio 3 software. All values were then normalized against the small RNA RNU6^34^.

#### Western blots

P2L MLFs were grown to confluence in 35 mm dishes and synchronized via temperature cycling. A paired plate of cells was handled similarly and recorded for bioluminescent traces as described above. Before recording, the media for all plates was changed into recording media (CO_2_). After treatment, cells were harvested and total protein was extracted by cell lysis by application of 2X Laemmli buffer supplemented with HALT Protease Inhibitor Cocktail (Fisher) and HALT Phosphatase Inhibitor Cocktail (Fisher). Proteins were denatured by boiling for 5 minutes. SDS-PAGE was performed to separate the proteins in the samples and the proteins were transferred to a polyvinylidene chloride (PVDC) membrane using a Bio-Rad Transfer-Blot Turbo (Bio-Rad). Membranes were blocked in Tris-HCl-buffered saline with 0.1% Tween-20 and 5% bovine serum albumin or skim milk for at least 30 minutes. Membranes were then incubated with p-p38 (Cell Signaling 9211S), p38 MAPK (Cell Signaling 9212S), p-ERK (Cell Signaling 9106S), ERK (Cell Signaling 9102S), p-eIF4E (Cell Signaling 9741S), eIF4E (Cell Signaling 9742S), or actin (Cell Signaling 3700S) primary antibodies at 1:1000 dilution for 1 hour before incubating with Goat anti-Rabbit, HRP conjugate (Invitrogen 31460) or Goat anti-Mouse, HRP conjugate (Invitrogen 32430) secondary antibodies at 1:5000 for 1 hour. Blots were visualized with ClarityMax ECL substrate (Bio-Rad) or Pierce ECL substrate (Thermo Scientific) and imaged in a Bio-Rad ChemiDoc MP (Bio-Rad). Images were quantified with the FIJI software^71^, and statistical analyses of quantifications were performed in Prism 10.

#### ELISA

P2L MLFs were grown to confluence in 96-well plates and kept at constant 37°C, 5% CO_2_. PRR agonists (Mouse TLR1-9 Agonist Kit, Invivogen tlrl-kit1mw) were diluted in DMEM supplemented with 10% (v/v) FBS to the following concentrations: Pam3CSK4 (0.3 μg/mL), PolyI:C (10 μg/mL), LPS (5 μg/mL), Flagellin (1 μg/mL), FSL-1 (0.1 μg/mL). Cells were stimulated by aspirating all cell culture media, adding the diluted agonists, and incubating at 37°C, 5% CO_2_ for 24 hours. The cell culture media was collected and stored at −20°C upon completion of the stimulation. IL-6 concentrations were quantified using the Mouse IL-6 ELISA MAX Set Standard kit (BioLegend 431301).

#### Mathematical modeling of the circadian clock

A custom script was written in Python to simulate the functional mechanism of a circadian clock. The script generates the simulation by continuously iterating upon the current state of the simulated clock according to a set of rules. Briefly, activated enhancer protein A produces the repressor protein B in a linear fashion. In turn, protein B inactivates protein A. At every iteration, a set percentage of protein B is degraded. To simulate the kinetics of protein binding, translation, and translocation, changes in the production of protein B or the inactivation of protein A are delayed according to a *production_change_rate* variable and a *repression_rate* variable, respectively. These variables, along with the protein B degradation rate, were optimized by iterative testing to produce oscillations with a 24-hour period. To model community-driven dynamics, a stochastic modeling step was added to the production and degradation of protein B, such that the production and degradation rates are multiplied by a Gaussian random variable with a mean of 1 and a standard deviation of 0.20. This randomized oscillator was then independently simulated for 3000 iterations, and the output was obtained by averaging the outputs of the 3000 simulations. The simulation script is released on Github (https://github.com/devmo24/Circadian-oscillator-simulation) under the GNU General Public License (GPL) v3.0.

#### Data presentation and statistical analysis

Detailed statistical reporting including sample size, p values, and the statistical test performed in every figure is provided in **Supplementary Table 2**. Briefly, all experiments were completed in triplicate or quadruplicate. Individual wells were considered distinct samples, and samples were measured once per timepoint. Luminescence recording traces are graphically grouped by experiment, such that if a PBS or bacterial control is shown on the same graph, it came from the same experiment. For the purposes of statistical analysis, measurements were assumed to be normally distributed with equal standard deviations across treatment groups. Where treatment groups are compared to a negative control treatment, statistical comparisons were calculated by a two-tailed unpaired t-test (for experiments with two treatment groups) or ANOVA with Dunnett’s multiple comparisons test (for experiments with more than two treatment groups).

Where comparisons are made across multiple treatment groups within a single experiment, statistical comparisons were calculated by ANOVA with Tukey’s multiple comparisons test. For the inhibitor effect compilation graph (Extended data Fig. 5E), which contained data from multiple experiments, inhibitors were compared only to their within-experiment (*Spn* + mock) groups using ANOVA with Sidak’s multiple comparisons test. Western blot quantifications were compared using Two-way ANOVA with Sidak’s multiple comparisons test. Effect sizes and degrees of freedom were calculated as recommended by Prism 10. All comparisons where p<0.05 were considered statistically significant.

## Supplementary Figures

**Extended data Fig. 1.**
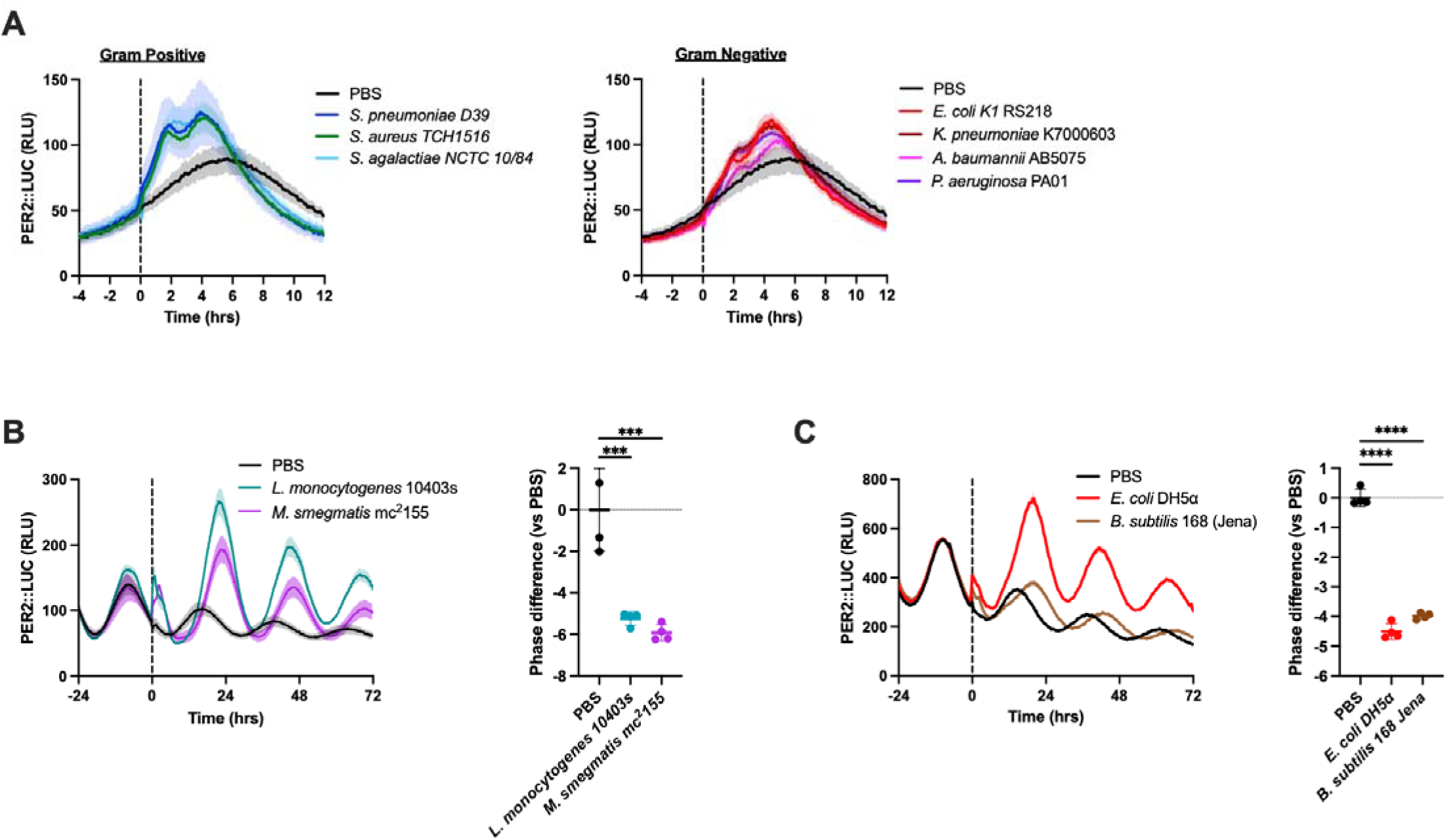
Exposure to additional bacterial species induces phase shifts. (**A**) A zoom in of −6 to 12-hr of the luminescence traces shown in Fig. 1A is shown to visualize acute changes in luminescence following HK bacteria exposure at time 0 (dashed line). Relative Luminescence Units (RLU). (**B-C**) P2L MLFs were stimulated with (B) *L. monocytogenes*, *M. smegmatis* or (C) *B. subtilis*, *E. coli* at time 0 (dashed line). Recording traces are shown as mean±S.E.; all other graphs are shown as mean±S.D. ***p<0.001; ****p<0.0001. *n* ≥ *3* for all groups, see Table S2 for all sample sizes and detailed statistical results.

**Extended data Fig. 2.**
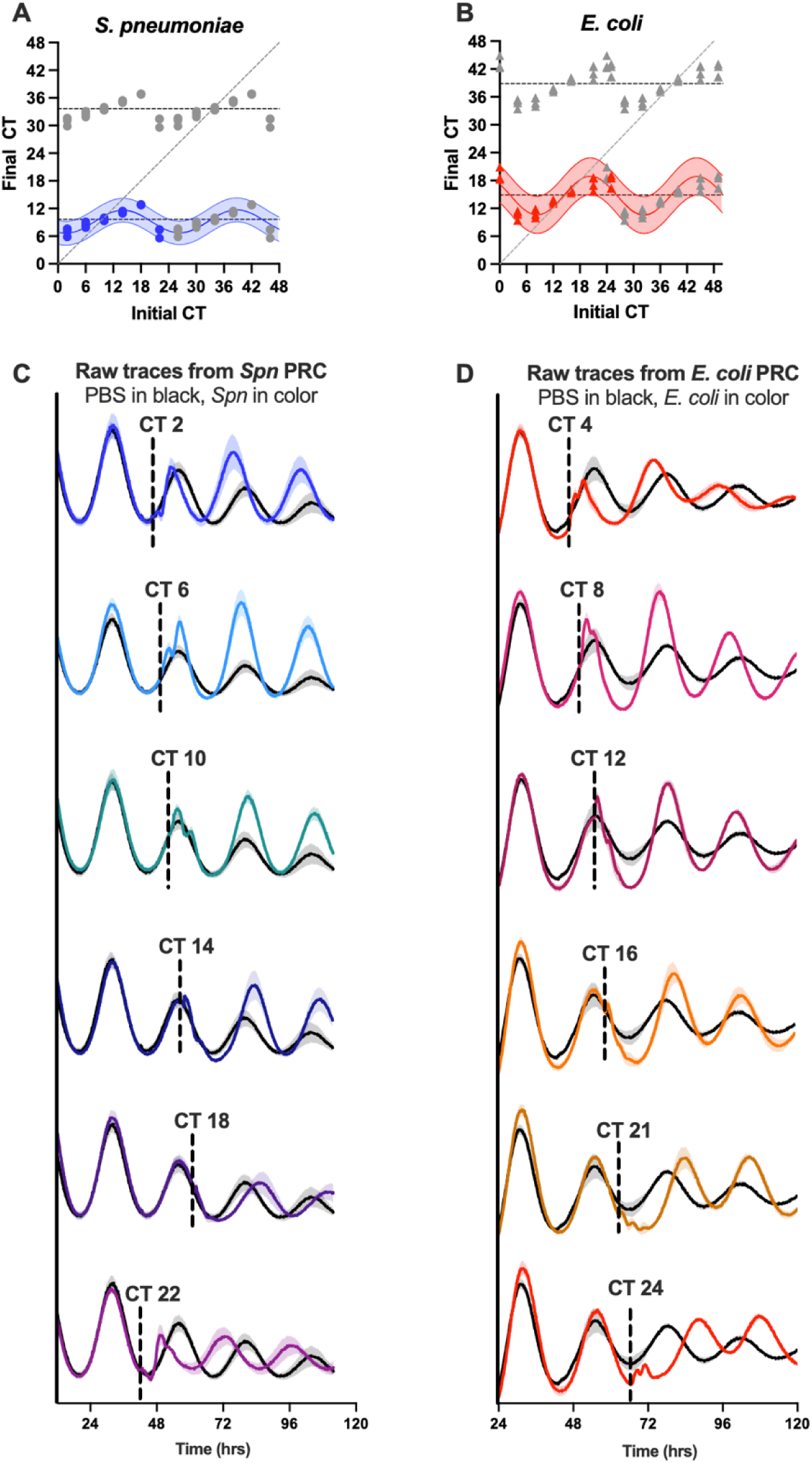
Raw data related to stimulations shown in Figure 3. (**A-B**) The data presented in Fig. 3C is presented here as a phase transition curve for *Spn* (A) and *E. coli* (B). The data is double-plotted along the horizontal axis and additionally double-plotted along the vertical axis, with a singular data plot shown in blue (for *Spn*) and red (for *E. coli*) representing the actual data collected. Duplicated data plots are depicted in gray. Initial circadian time (CT) indicates the circadian phase at the time of stimulation; final CT indicates the phase post-stimulation. Horizontal dashed lines indicate the average final CT across all stimulations. The diagonal dashed line is a line of identity and has a slope of 1. An oscillation of best fit is shown as a colored line on the graph. Shading around the colored line indicates the 95% confidence interval of the oscillation. (**C-D**) Recording traces for the phase response curves (shown in Fig. 3C) for *Spn* (C) and *E. coli* (D). Colored traces in (C-D) represent the mean signal of *n* ≥ *3* samples, shading indicates S.E. Bacteria were added at specific phases (labeled in circadian time, CT) as indicated by the dotted line.

**Extended data Fig. 3.**
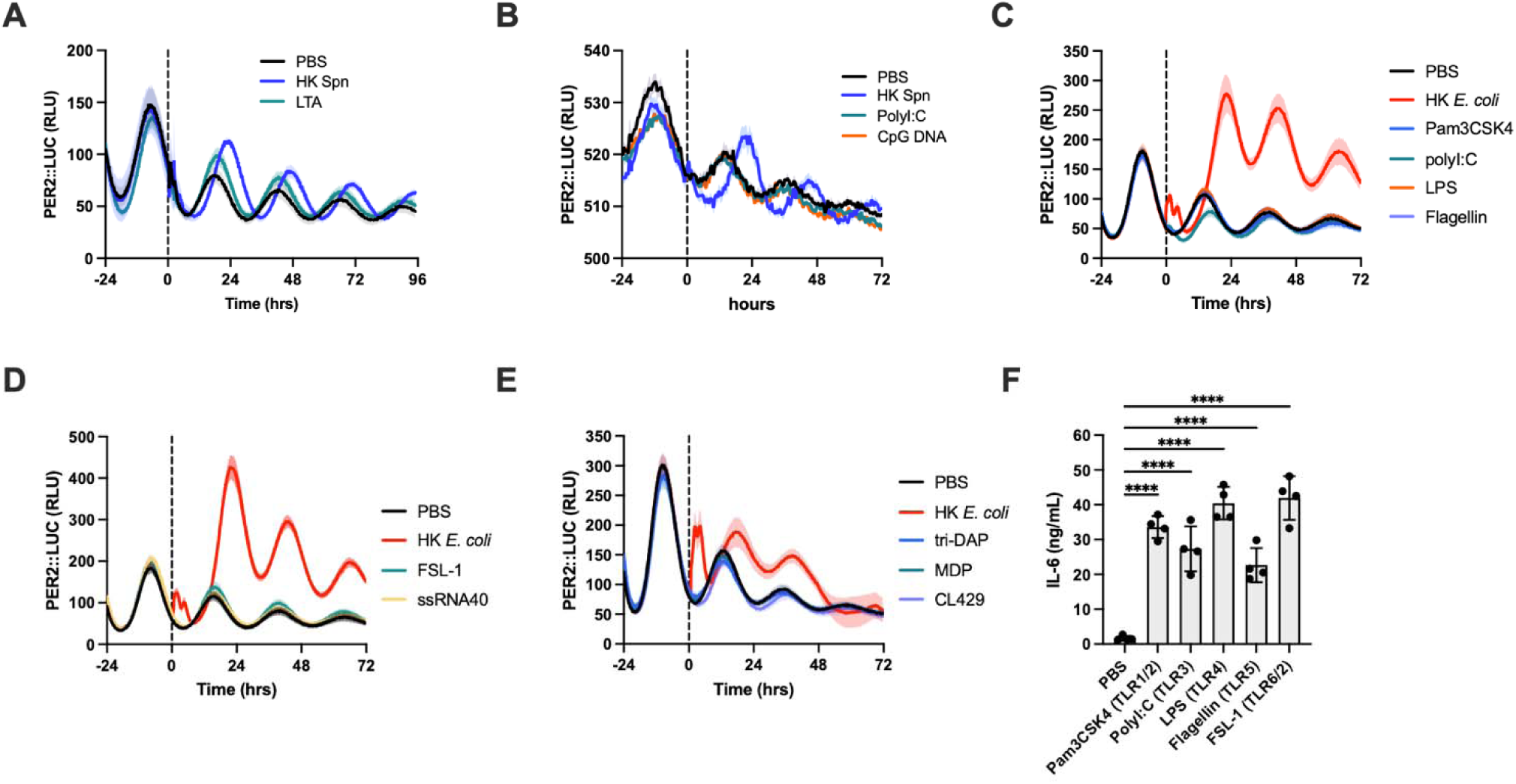
Raw data related to stimulations shown in Figure 4. (**A-E**) Recording traces for the data presented in Fig. 4A are shown. An additional experimental replicate with LTA stimulation is shown in (A). (**F**) Unsynchronized P2L MLFs were stimulated with various pattern recognition receptor (PRR) agonists and analyzed for IL-6 secretion. Recording traces are shown as mean±S.E.; (F) is shown as mean±S.D. **** = p<0.0001. *n* ≥ *3* for all groups, see Table S2 for all sample sizes and detailed statistical results.

**Extended data Fig. 4.**
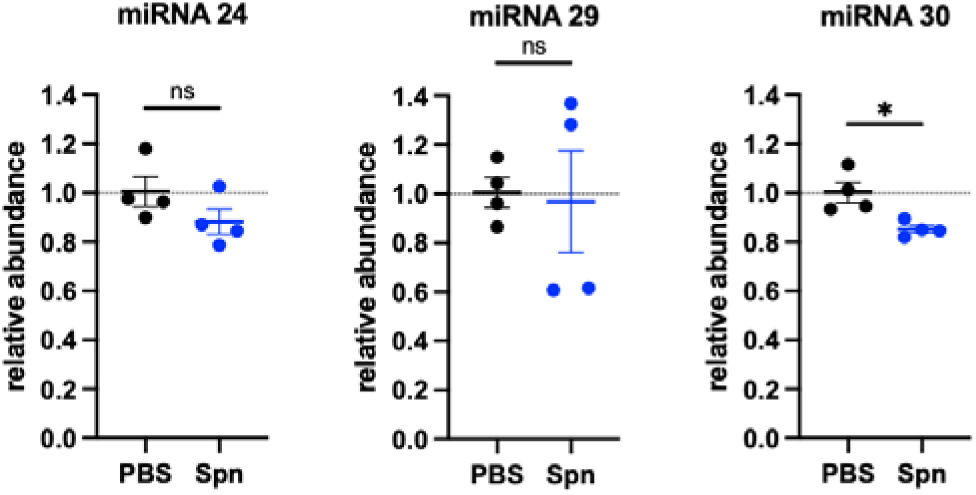
miRNA abundance is unchanged following microbial exposure. miRNA abundance in P2L MLFs was quantified by qRT-PCR at 4-hr post stimulation with HK *Spn* or PBS. Graphs are shown as mean±S.D. * = p<0.05; ns = not significant. *n = 4* for all groups, see Table S2 for all sample sizes and detailed statistical results.

**Extended data Fig. 5.**
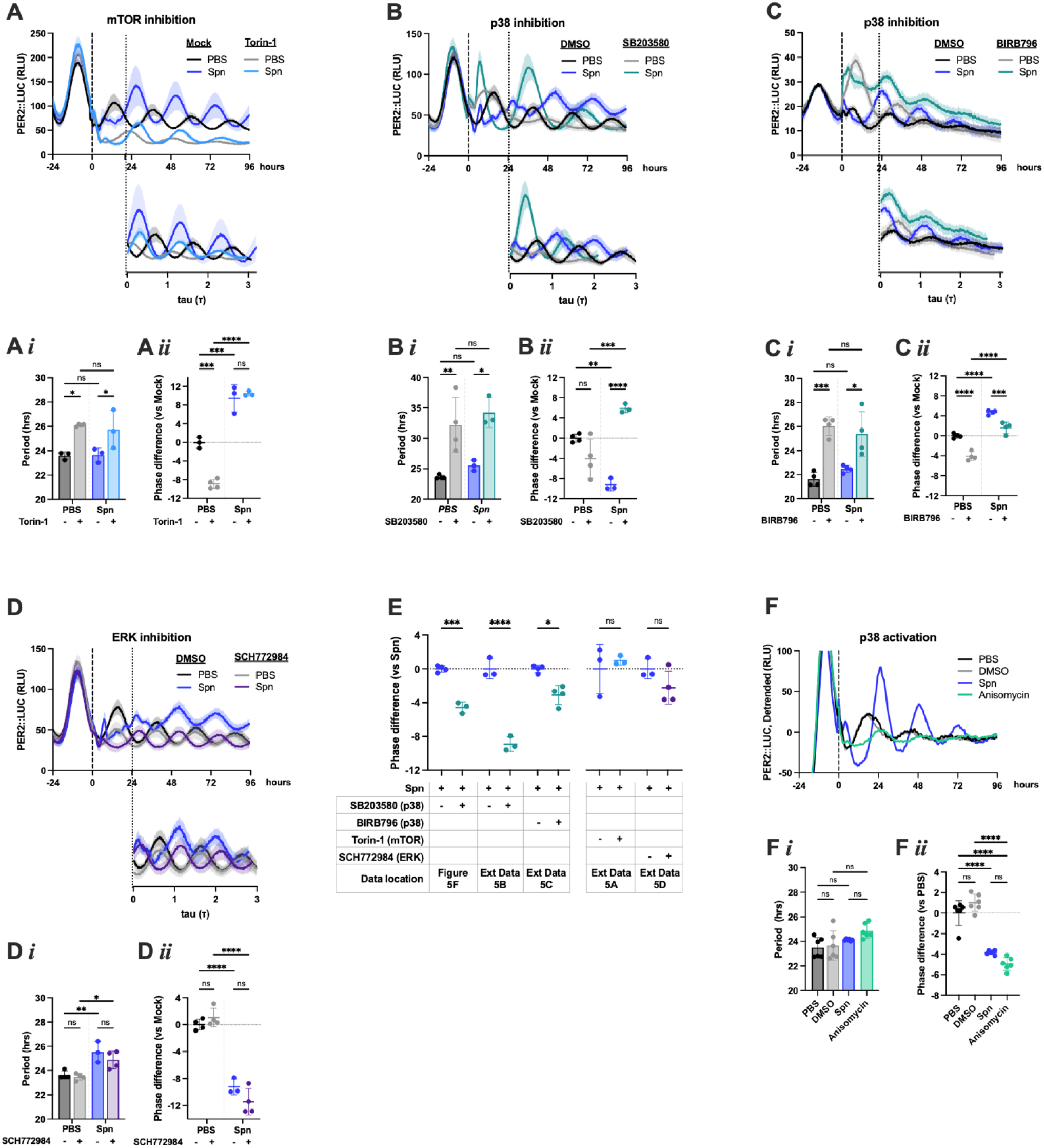
p38 MAPK inhibition uniquely modulates microbe-induced phase shifts. Additional data demonstrating responses of P2L cells stimulated with PBS or HK *Spn* in the presence of select pharmacologic inhibitors. Recording traces are shown as relative light units (RLU) graphed in hours and, below, in modulo-*tau* format normalized based on period (τ) for each group. Modulo-*tau* graphs begin when the oscillations stabilize post-stimulation, indicated by the dotted line connecting the two graphs. Period was determined from the raw RLU traces, and used to generate modulo-*tau* graphs. Groups with a longer period end earlier in modulo-*tau* format as fewer circadian cycles were completed within 96 hr of recording. (**A**) Effect of Torin-1 on P2L oscillations (top), period (*i*), and differences in phase (*ii*) compared to PBS controls. (**B**) Effect of SB203580, an inhibitor of p38 MAPK, on P2L oscillations (top), period (*i*), and differences in phase (*ii*) compared to PBS controls. Data is an independent experimental repeat of Fig. 5F. (**C**) Effect of BIRB796, an inhibitor of p38 MAPK, on P2L oscillations (top), period (*i*), and differences in phase (*ii*) compared to PBS controls. (**D**) Effect of SCH772984, an inhibitor of ERK MAPK, on P2L oscillations (top), period (*i*), and differences in phase (*ii*) compared to PBS controls. **(E)** Phase difference between *Spn* + inhibitor and *Spn* + mock control. Data is from Fig. 5F and Extended data 5A-D (this figure). (**F**) P2L MLFs were stimulated with HK *Spn* or anisomycin, with PBS or DMSO as controls. Shown is the effect of the stimulation on P2L oscillations (top), period (*i*), and differences in phase (*ii*) compared to PBS controls. Recording traces are shown as mean±S.E.; all other graphs are shown as mean±S.D. * = p<0.05; ** = p<0.01; *** = p<0.001; **** = p<0.0001; ns = not significant. *n* ≥ *3* for all groups, see Table S2 for all sample sizes and detailed statistical results.

**Extended data Fig. 6.**
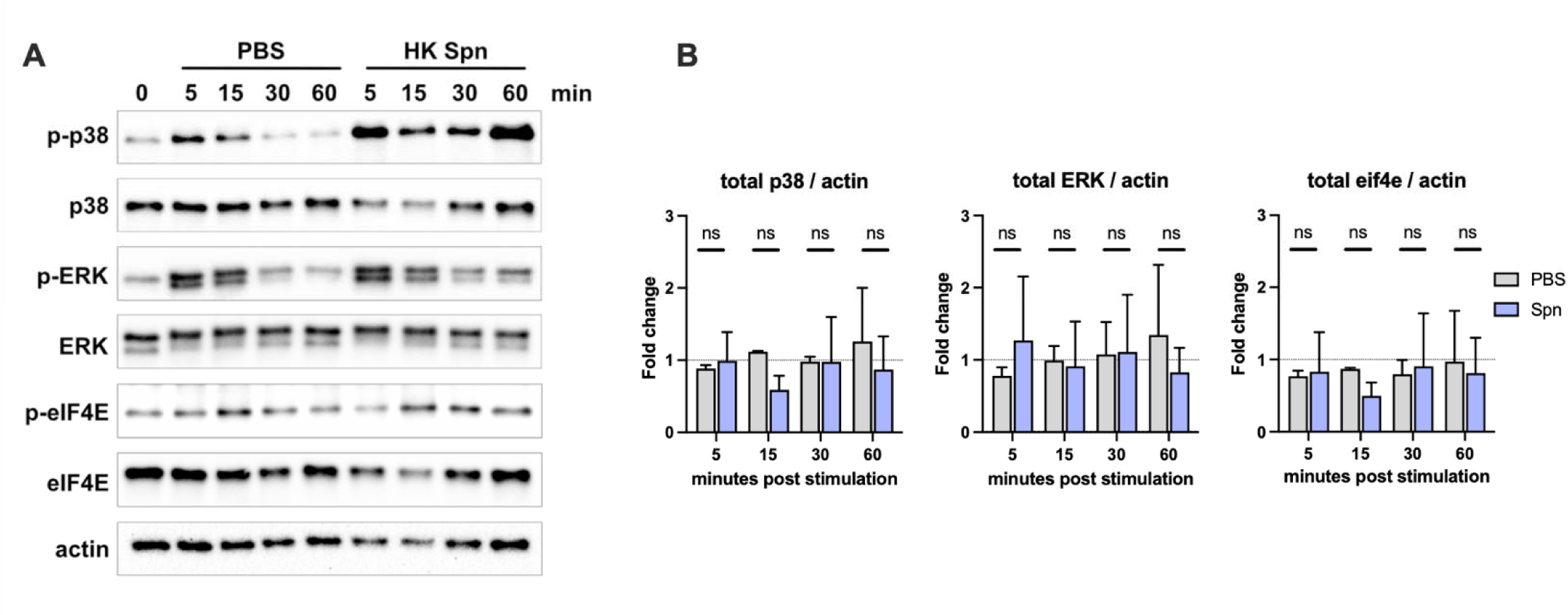
Additional replicates for Western blot analyses shown in Figure 5. (**A**) An additional replicate of cell lysates was collected and blotted for total and phosphorylated p38, ERK, and eIF4E. Actin is a loading control. (**B**) Densitometry analysis of Western blots shown in Fig. 5G and (A). The relative abundance of protein was calculated at each timepoint as [total protein] / [actin]. The fold change in activation was calculated by comparing each timepoint to the 0 minute sample on the same blot.

**Extended data Fig. 7.**
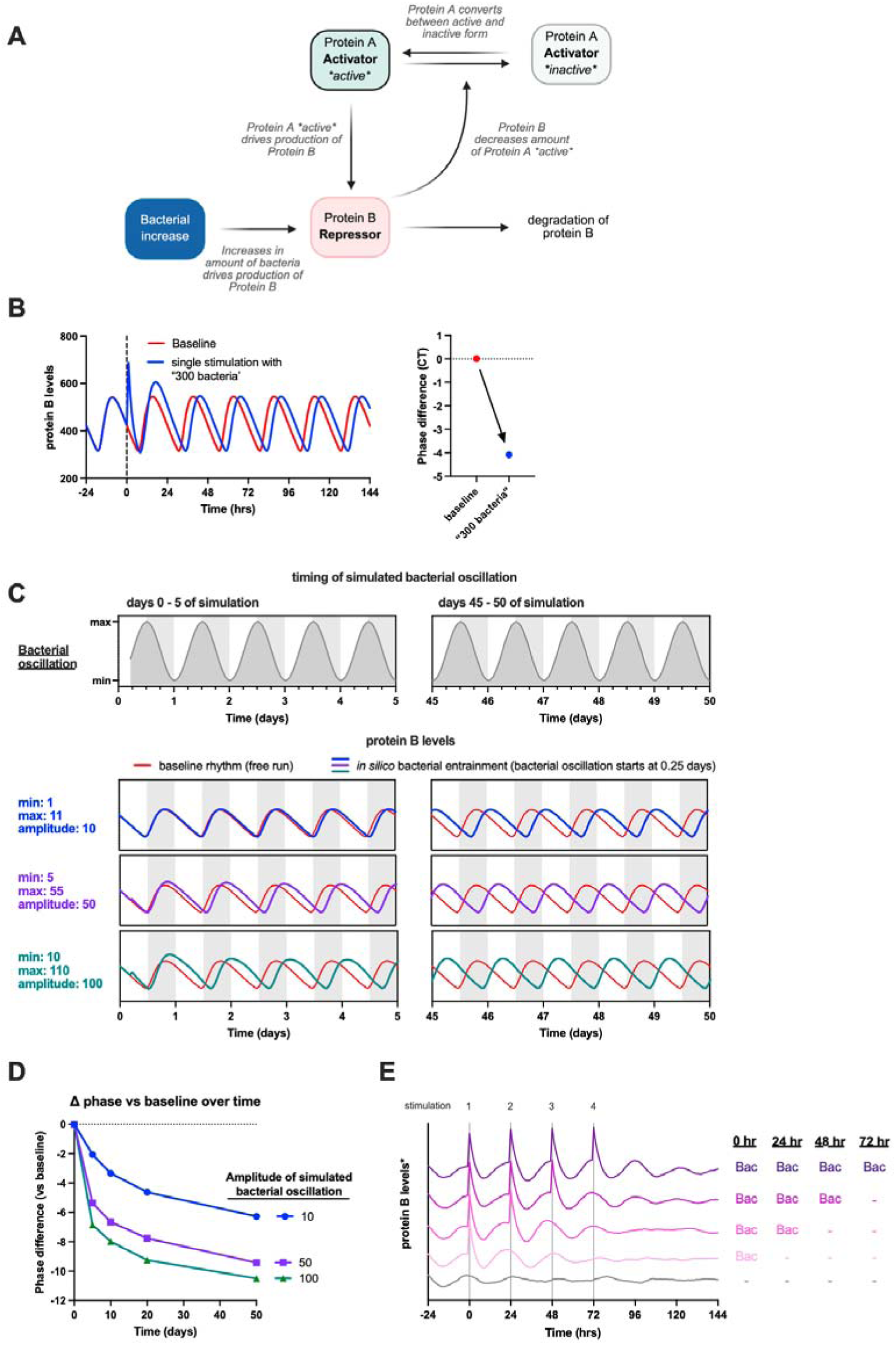
A mathematical model predicts environmental bacteria are a circadian entrainment cue. (**A**) A diagram depicting the interactions used in the simulation model. In subsequent graphs, the levels of “Protein B (Repressor)” are shown. (**B**) Levels of Protein B are modeled at baseline (red) or following addition of “300 bacteria” (blue) added at time 0 (vertical dashed line). An acute increase in Protein B occurs immediately and drives a change in the resulting phase of the oscillation. The phase difference (right) of the resulting simulated oscillation is calculated in circadian hours compared to the control (baseline). (**C-D**) The effect of daily oscillations in the amount of bacteria was modeled *in silico* to simulate how protein B responds over the course of 50 days. All simulations modeled a 10-fold increase in bacteria over a minimum, baseline amount. Three minimum baseline amounts were used (1, 5, or 10 bacteria) which leads to amplitudes of 10, 50, or 100 in the bacterial oscillations. The grey bands denote the 12 hour period when bacterial amount is greatest, and the white bands indicate when bacterial amount is lowest. **(C)** Days 0 - 5 of the simulation are shown on the left, and days 45 - 50 are shown on the right. Bacterial oscillations began at 0.25 days. All conditions lead to changes in the phase of Protein B, with higher amplitude oscillations exerting a stronger effect and driving more rapid adjustments. **(D)** Difference in phase of Protein B between baseline conditions (no bacteria present) and in the presence of bacterial oscillations was calculated at multiple timepoints along the stimulation. (**E**) Randomness in the production and degradation rate of Protein B is simulated at every simulation step, and measurements are recorded as the average of 3000 independent simulations. The effect of rhythmic addition of bacteria (time 0-, 24-, 48-, and 72-hrs; solid grey lines) on Protein B levels is shown. * in the vertical axis label indicates simulated traces were distributed for visual clarity.

## Tables

**Table S1. Bacterial strains and growth conditions**

**Table S2. Detailed statistical reporting including sample sizes, statistical comparisons and exact *p* values.**

## Use of AI

The large language model ChatGPT (OpenAI) was used to reduce the word count of original text written by the authors by submitting individual sentences or paragraphs with instructions to “say this in less words” or “point out unnecessary redundancy.” Individual changes to the text were then manually incorporated into the manuscript by the authors as appropriate.

## Notes

### Competing Interest Statement

The authors have declared no competing interest.

